# Spontaneous and coordinated Ca^2+^ activity of cochlear sensory and non-sensory cells drives the maturation of OHC afferent innervation

**DOI:** 10.1101/317172

**Authors:** Federico Ceriani, Aenea Hendry, Jing-Yi Jeng, Stuart L. Johnson, Jennifer Olt, Matthew C. Holley, Fabio Mammano, Corné J. Kros, Dwayne D. Simmons, Walter Marcotti

## Abstract

Outer hair cells (OHCs) are highly specialized sensory cells conferring the fine tuning and high sensitivity of the mammalian cochlea to acoustic stimuli. Here, by genetically manipulating spontaneous Ca^2+^ signalling *in vivo*, through a period of early postnatal development, we find that the refinement of OHC afferent innervation is regulated by complementary spontaneous Ca^2+^ signals originating in OHCs and non-sensory cells. OHCs fire spontaneous Ca^2+^ spikes during a narrow period of immature development. Simultaneously, waves of Ca^2+^ activity in the non-sensory greater epithelial ridge act, via ATP-induced activation of P2X receptors, to synchronize OHC firing, resulting in the refinement of their afferent innervation. In the absence of connexin channels Ca^2+^ waves are impaired, leading to a reduction in the number of ribbon synapses and afferent fibres on OHCs. We propose that the correct maturation of the afferent connectivity in OHCs requires experience-independent Ca^2+^ signals from sensory and non-sensory cells.

## Introduction

Mammalian hearing depends upon two specialized sensory receptors in the organ of Corti, the inner and outer hair cells, and their afferent and efferent neuronal connections, the differentiation and maturation of which requires precise timing and coordination between genetic programmes and physiological activity (**Corns et al., 2014; Delacroix and Malgrange, 2015**). Inner hair cells (IHCs) are the primary sensory receptors and they relay sound information to spiral ganglion afferent neurons via the release of glutamate from vesicles tethered to presynaptic ribbons. By contrast, the role of outer hair cells (OHCs) is to contribute to the large functional dynamic range of the mammalian cochlea and to enhance the sensitivity and the frequency tuning within the cochlear partition (**Dallos, 1992**). Adult OHCs are primarily innervated by the cholinergic medial olivocochlear neurons (**Liberman, 1980; Maison et al., 2003**), the role of which is to modulate mechanical amplification in the adult cochlea (**Guinan, 1996**). However, OHCs are also innervated by Type II afferent fibres that, unlike the Type I fibres contacting IHCs, which encode sound timing, intensity and frequency, are activated solely by acoustic trauma (**Flores et al., 2015; Liu et al., 2015**). In most altricial rodents OHCs only begin to acquire the innervation pattern present in the mature cochlea towards the end of the first and the start of the second postnatal week (**Simmons, 1994; Simmons et al., 1996**). However, the molecular mechanisms responsible for the correct afferent innervation of OHCs remain poorly understood.

The refinement of sensory circuits during development is normally influenced by periods of experience-independent action potential (AP) activity before the onset of function (**Katz et al., 1996; Blanckenship et al., 2010**). Calcium-dependent APs have been shown to occur spontaneously in immature IHCs (**Johnson et al., 2011; 2017**) but not in OHCs (**Marcotti and Kros, 1999; Oliver et al., 1997; Weisz et al., 2012**). One study reported spontaneous APs in OHCs of wild-type and otoferlin mutant mice, but mostly using elevated extracellular Ca^2+^ and high intracellular EGTA (**Beurg et al., 2008**).

We found that during a narrow, critical period of postnatal development, OHCs fire spontaneous Ca^2+^ spikes immediately preceding their functional maturation (~P7-P8). These spikes can be modulated by ATP-induced Ca^2+^-waves travelling amongst non-sensory cells to synchronize the activity of nearby OHCs. *In vivo* alteration of the Ca^2+^ dependent activity in non-sensory cells prevented the maturation of the OHC afferent innervation. We propose that precisely modulated spontaneous Ca^2+^ signals between OHCs and non-sensory cells are necessary for the correct maturation of the neuronal connectivity to OHCs.

## Results

The functional development of OHCs was studied primarily in the apical third of the mouse cochlea, corresponding to a frequency range in the adult mouse of ~6-12 kHz (**Müller et al., 2005**) (**Figure 1A**). For comparison, some recordings were also made from the basal coil of the cochlea through the frequency range of ~25-45 kHz (**Figure 1A**). Spontaneous activity in immature OHCs and its modulation by non-sensory cells in the greater (GER) and lesser (LER) epithelial ridges (**Figure 1B**) were recorded from cochleae bathed in a perilymph-like extracellular solution (1.3 mM Ca^2+^ and 5.8 mM K^+^) (**Wangemann and Schacht, 1996**) either near body temperature or at room temperature. The stereociliary bundles of hair cells (**Figure 1B**) are normally bathed in endolymph, which contains ~150 mM K^+^ and ~20 μM Ca^2+^ in the mature cochlea (**Wangemann and Schacht, 1996**; **Bosher & Warren, 1978**). However, during the first few days after birth, endolymph has a similar ionic composition to that of the perilymph (**Wangemann and Schacht, 1996**).

**Figure 1.**
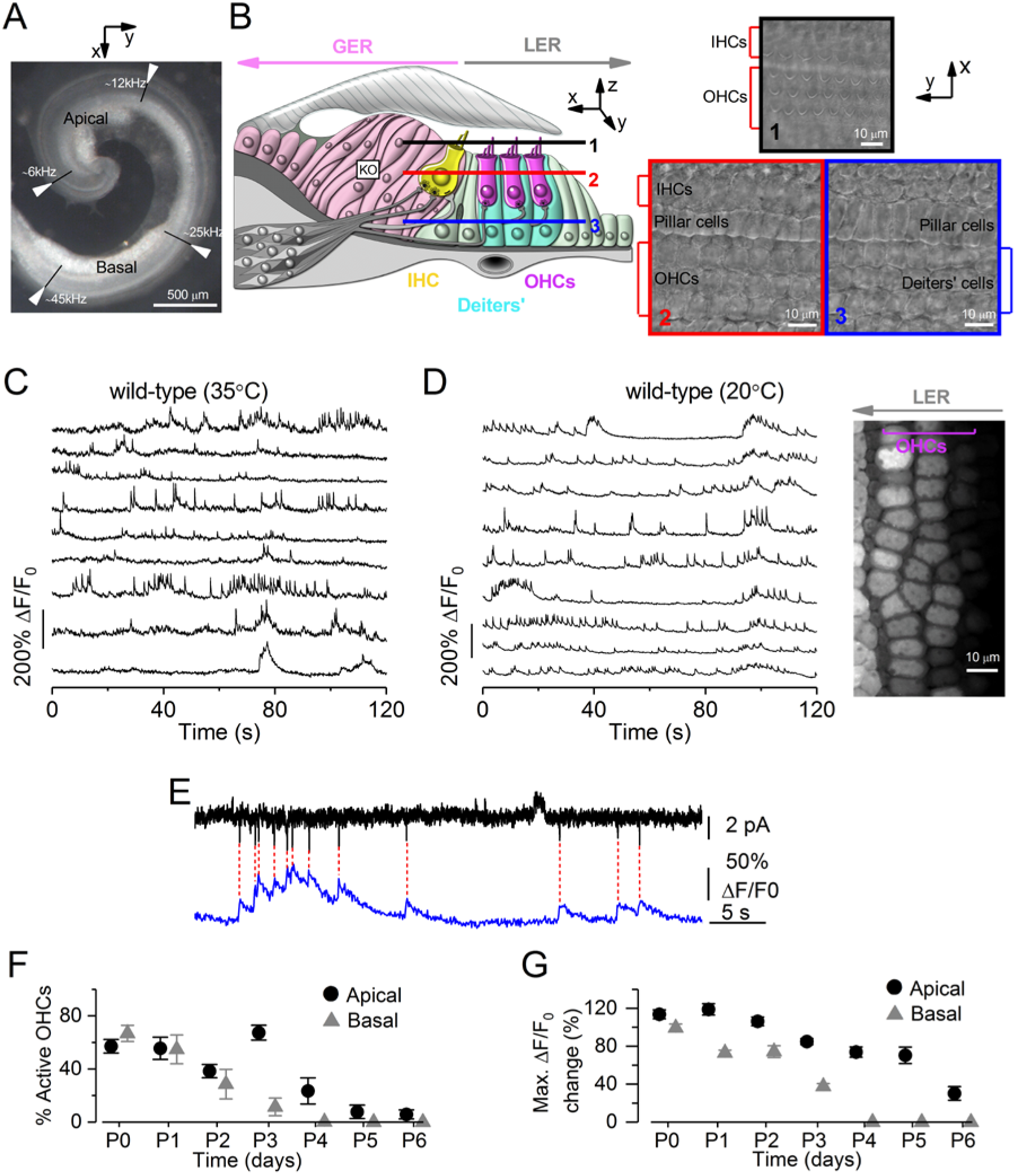
Early postnatal OHCs show spontaneous Ca^2+^ spikes. (**A**) Image of the mouse cochlea at P2 highlighting the apical and basal regions used for the experiments. The apical and basal regions were at a fractional distance along the coil of approximately 8% to 32% (corresponding to a frequency range in the mature mouse of ~6-12 kHz) and 55% to 80% (~25-45 kHz) from the apex, respectively. (**B**) Diagram (left) showing a cross-section of an early postnatal organ of Corti. OHCs: outer hair cells; IHCs: inner hair cells; GER: greater epithelial ridge, which includes non-sensory cells such as the inner phalangeal cells (surrounding the IHCs) and tightly packed tall columnar cells forming the Kölliker’s organ (KO); LER: lesser epithelial ridge. Right panels show DIC images of the cochlea at the level of the hair bundle (top) and both OHCs and non-sensory Deiters’ cells in the LER region (bottom). (**C,D**) Representative ΔF/F0 traces from 9 apical OHCs of a P2 wild-type mouse (from the image in the right panel) recorded at body (**C**) and room (**D**) temperature. Traces are computed as pixel averages of regions of interest centred on OHCs. Calcium spikes are evident in OHCs. In this and the following panels, the right image provides a visual representation of the spontaneous activity over the entire duration of the recordings (120 s), and was obtained by averaging 4000 frames of raw data. (**E**) Simultaneous cell-attached patch clamp recording (top) and Ca^2+^ imaging (bottom) obtained from a P1 OHC from wild-type mice at room temperature. The intracellular Ca^2+^ level in OHCs increased rapidly but decayed with a relatively long fluorescence decay time constant (~300 ms) (**Ceriani et al., 2016**). (**F**) Percentage of apical and basal OHCs showing spontaneous Ca^2+^ spikes at near body temperature and as a function of postnatal age. Note that these values likely represent an underestimation of the fraction of active OHCs, since the three-dimensional structure of the cochlea and the optical sectioning capability of 2-photon microscopy make it difficult to be in the optimal focal conditions for the simultaneous recording of all three rows of OHCs. Moreover, active OHCs were those showing activity within the 2 minute recording time. Number of total OHCs and recordings from left to right were: apical cochlea 384 and 8; 338 and 7; 441 and 10; 497 and 9; 466 and 10; 453 and 11; 254 and 6; basal cochlea 408 and 9; 528 and 11; 460 and 10; 380 and 7; 676 and 12; 143 and 3; 133 and 4. (**G**) Maximum ΔF/F0 changes in apical and basal active OHCs as a function of postnatal age. Number of active OHCs were: apical cochlea 218; 189; 82; 336; 94; 39; 6; basal cochlea 273; 303; 124; 46; 0; 0; 0. These OHCs came from the dataset in (**F**).

### Calcium-dependent spikes in OHCs occur spontaneously during a narrow period of development

Spontaneous, rapid Ca^2+^ transients were recorded from OHCs maintained at near body (~35°C: **Figure 1C**, **Movie 1**) and room temperature (~20°C: **Figure 1D**, **Movie 3, top panel**) in acutely dissected cochleae from new-born mice loaded with the Ca^2+^ indicator Fluo-4). Similar Ca^2+^ transients were observed in OHCs in 37 separate recordings from 13 different mice. By combining cell-attached patch clamp recordings and Ca^2+^ imaging, we confirmed that the Ca^2+^ signals represent the optical readout of OHC firing activity, with bursts of APs causing large increases in the OHC Ca^2+^ level (**Figure 1E**). Although Ca^2+^ spikes were present in OHCs along the entire cochlea at birth, the number of cells showing this activity decreased over time, with basal OHCs being the first to stop at around P4 and apical cells stopping a couple of days later (**Figure 1F**). This correlated with a decrease in the maximum Ca^2+^-related changes in fluorescence intensity (Δ*F*/*F*_0_) for Ca^2+^ measured from active OHCs (**Figure `1G**), which could be due to a progressive reduction of AP bursts with age.

Calcium transients were abolished in Ca^2+^ free solution (**Figure 2A,B**, **Movie 2**) and were absent in OHCs lacking Ca_V_1.3 Ca^2+^ channel subunit (**Figure 2C; Movie 3, bottom panel**), which is the main Ca^2+^ channel subunit expressed in hair cells (**Platzer et al., 2000**). These results, together with the finding that Ca^2+^ spikes were not prevented when blocking Ca^2+^ release from intracellular stores (**Figure 2D**), indicate their dependence on extracellular Ca^2+^.

**Figure 2.**
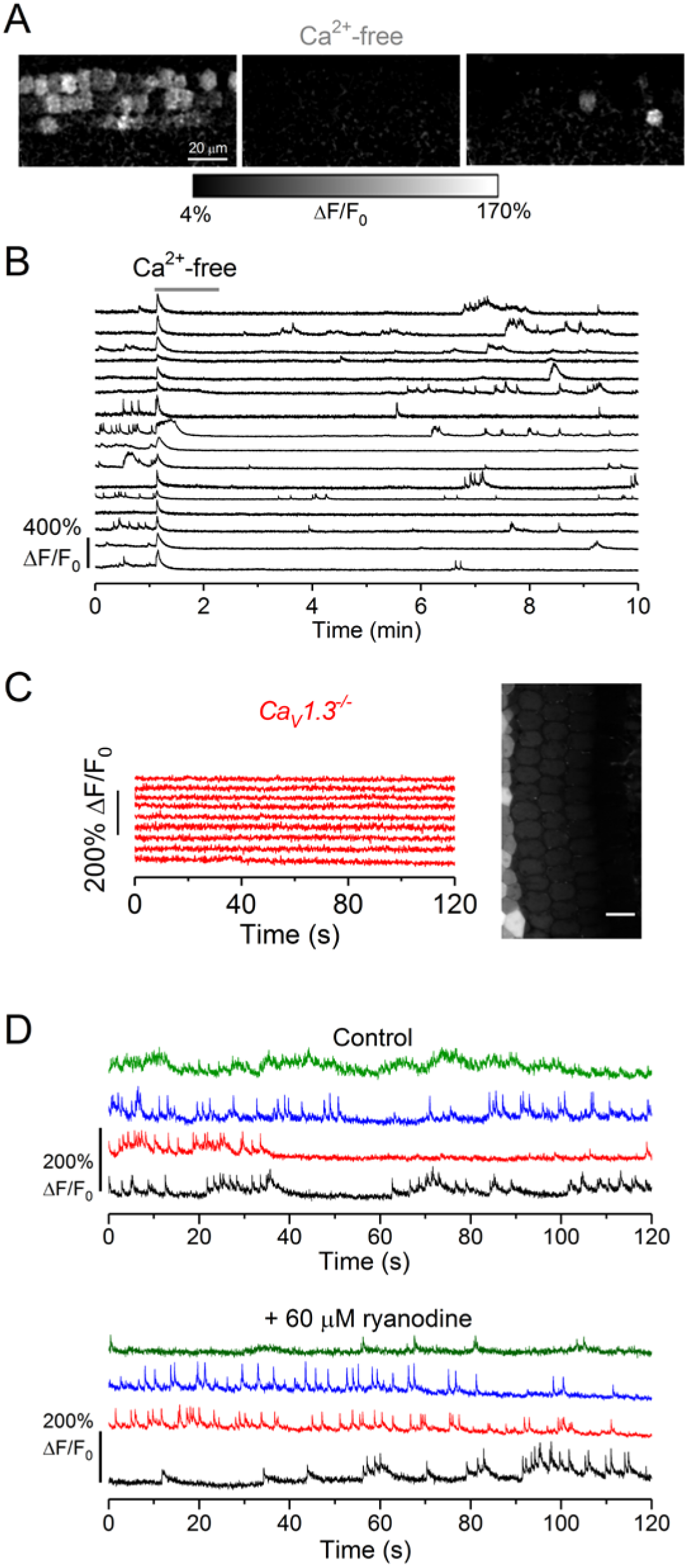
Spontaneous Ca^2+^ spikes in OHCs are Ca^2+^ dependent. (**A**) Relative fluo-4 fluorescence changes (ΔF/F_0_) before (left) during (middle) and after (right) the application of a Ca^2+^-free extracellular solution. Images were obtained as maximal back projections (500 frames, 16.5 s). (*B*) Representative ΔF/F_0_ traces from 16 OHCs (from the images in panel *A*) during the local application of the Ca^2+^-free extracellular solution with the Picospritzer. Traces are computed as pixel averages of regions of interest centred on OHCs. Ca^2+^ spikes are evident in OHCs before the application of the Ca^2+^-free extracellular solution. Note that the slow re-appearance of the Ca^2+^ spikes was due to the fact that the control solution containing 1.3 mM Ca^2+^ was bath perfused. A total of five recordings were performed from four cochleae (2 mice age P2). Recordings are at RT. (**C**) Representative ΔF/F0 traces from 9 apical OHCs of a P2 *Ca_V_1.3^-/-^* mouse (selected from the images in the right panel) recorded at room temperature. Calcium traces were computed as in **Figure 1C**. (**D**) Representative dF/F_0_ traces from the same four P2 OHCs (colour coded) before (top panel) and during (bottom panel) the application of 60 μM ryanodine, which blocks the Ca^2+^ efflux from the stores (**Meissner, 1986**). Note that ryanodine was applied for at least 30 minutes before re-imaging OHC activity. A total of 16 recordings were performed from 11 cochleae (7 mice age P2-P3). Recordings are at RT.

### Calcium-waves from non-sensory cells coordinate OHC electrical activity

Hair cells are embedded in a matrix of non-sensory, epithelial supporting cells (**Figure 1B**). The inner phalangeal cells surrounding the IHCs and the tightly packed columnar cells that form the Kölliker’s organ, are part of the GER and they show spontaneous inward currents (**Tritsch et al., 2007**). This spontaneous activity is initiated by extracellular ATP, which is released via an extensive network of connexin hemichannels in non-sensory cells, activating purinergic autoreceptors on the same cells, causing an increase in intracellular Ca^2+^. It leads to spatially and temporally coordinated Ca^2+^ waves that are propagated across the epithelium (**Tritsch et al., 2007**). Calcium waves can also be triggered by the application of ATP to non-sensory cells surrounding the OHCs in the LER (e.g Deiters’ cells: see **Figure 1B**) but these waves are thought not to occur spontaneously (**Tritsch et al., 2007**). Therefore, we investigated whether spontaneous Ca^2+^ waves originating in the GER of the developing mouse cochlea (**Figure 3A**, red arrow) can influence the electrical activity in OHCs. We found that uncorrelated spontaneous bursts of Ca^2+^ activity in neighbouring OHCs became synchronized during large Ca^2+^ waves (**Figure 3B**, **Movie 4**) but remained unchanged during the smaller Ca^2+^ signals (**Figure 3C, Movie 5**). We quantified the average correlation coefficient (*r_s_*: see **Materials and Methods**) between every pair of OHCs in the field of view (64 ± 5 OHCs, 7 cochleae, 6 mice) during a time window of 400 frames (13.2 s) centred to the maximum intensity of the spontaneous Ca^2+^ signal occurring in the GER (grey area in **Figure 3B,C**, right panels). The average cross-correlation coefficients in nearby OHCs increased with the longitudinal (i.e. along the tonotopic axis) extension of Ca^2+^ waves in the GER (**Figure 3D**). While half of the smaller Ca^2+^ waves that spread over less than 75 μm (28 out of 56) had no significant effect on the correlation, all 19 of the larger waves synchronized the activity in OHCs (**Figure 3D**). However, the *r_s_* was independent of the amplitude (Δ*F/F_0_*) of the Ca^2+^ signal (**Figure 3E**) measured as a pixel average over the entire spread of the Ca^2+^ wave. Therefore, the coordination of the electrical activity between nearby OHCs was dependent on the lateral spread, but not the amplitude, of the Ca^2+^ waves.

**Figure 3.**
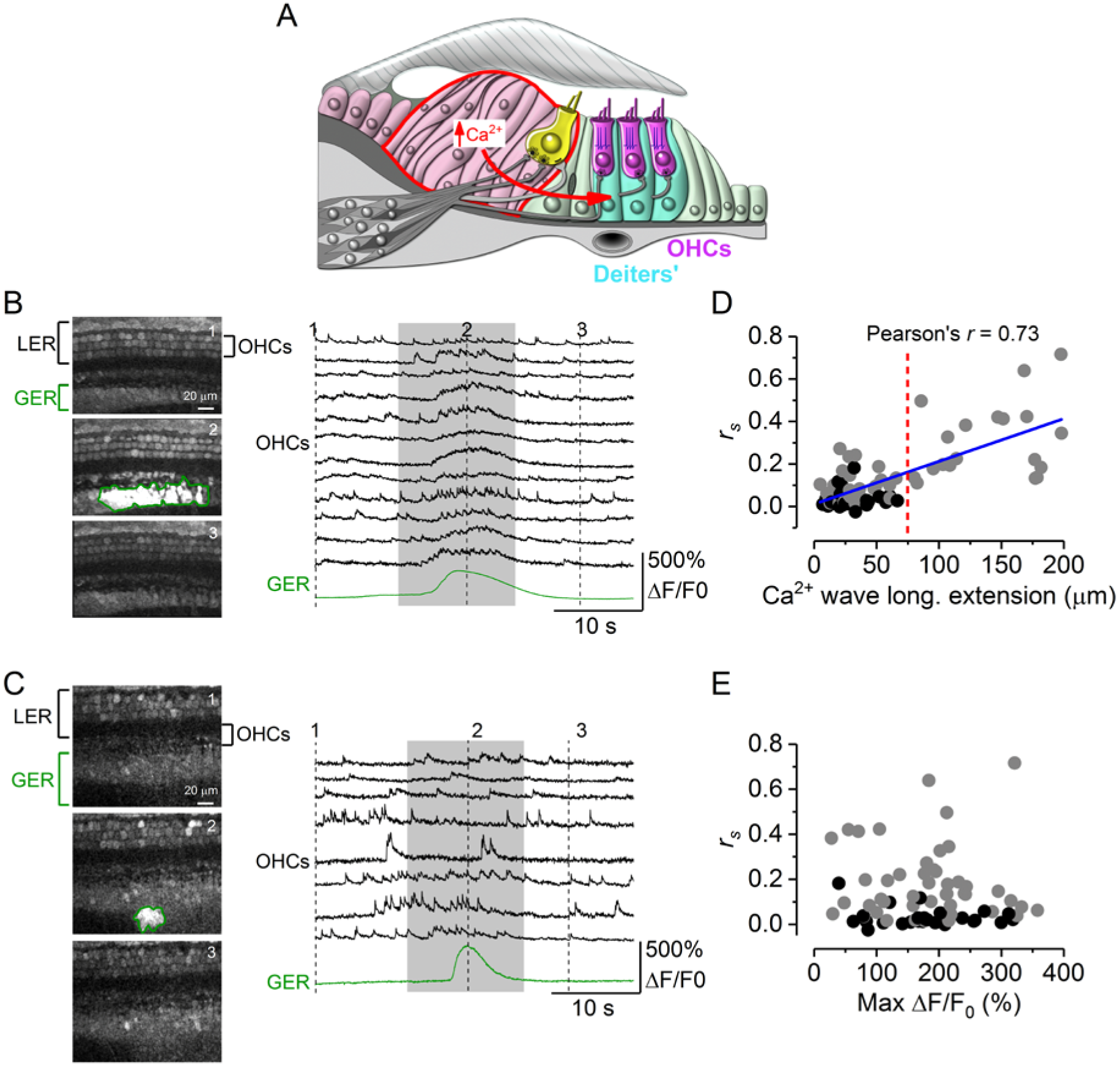
Calcium waves from the GER modulate OHCs spontaneous Ca^2+^-signalling. (**A**) Diagram showing a cross-section of an immature organ of Corti. Spontaneous Ca^2+^ waves (red arrow) are generated in the non-sensory cells present in the greater epithelial ridge (GER: red line). (**B** and **C**) Three representative images (left panels) obtained before, during and after the spontaneous appearance of a wide (**B**) and a narrow (**C**) Ca^2+^ wave in the GER of the apical coils of P2 wild-type mice. Right panels show representative ΔF/F0 traces from 12 (**B**) and 8 (**C**) OHCs (black traces) and those originating from the Ca^2+^ wave in the GER (green traces). The grey-shaded areas highlight the time window used for correlation analysis (see below). Recordings are at 31°C. (**D** and **E**) Average Spearman’s rank correlation coefficient (*r*_s_: see **Materials and Methods**) between the OHC activity as a function of the longitudinal extension (**D**) or the maximum ΔF/F0 changes (**E**) of spontaneous Ca^2+^ wave in the GER from the apical coil of P1-P2 mouse cochleae. The average length of the apical coil segments used for these experiments was 188 ± 4 μm (see **Materials and Methods**). Gray dots represent Ca^2+^ waves that are associated with a significant increase in OHC correlation, while black dots represent events during which OHC correlation didn’t increase significantly. Waves travelling more than 75 um in the longitudinal direction (red dashed line) always triggered a significant increase in OHC synchronization. Solid line in panel (**D**) represents a linear fit to the data. The slope was significantly different from zero (*P*<0.0001).

We then sought to identify how spontaneous Ca^2+^ activity from the non-sensory cells of the GER coupled to Ca^2+^ signalling in OHCs. Patch clamp recordings from Deiters’ cells, which surround the OHCs in the LER (**Figure 3A**), revealed spontaneous inward currents similar to those measured in non-sensory cells of the GER (**Tritsch et al., 2007**). During these spontaneous currents, Deiters’ cells depolarized by 16.7 ± 0.5 mV (range 5.4-43.5 mV, 339 events, *n* = 9) from an average resting membrane potential of −70.8 ± 1.6 mV (*n* = 9) (**Figure 4A,B**). Simultaneous recordings showed that the inward currents in the Deiters’ cells were synchronized with Ca^2+^ waves in the GER in both wild-type (**Supplementary Figure 1A**) and *Ca_V_1.3^-/-^* mice (**Figure 4C**). The absence of Ca^2+^ signals in OHCs from *Ca_V_1.3^-/-^* mice made it easier to see that the Ca^2+^ waves originating in the GER were able to travel to the LER and propagate through Deiters’ cells (**Figure 4D; Supplementary Figure 1B**). In order to test whether Deiters’ cells mediate signal transfer from the GER to the OHCs, we analysed Ca^2+^ signals after removal of Deiters’ cells (**Figure 5C**), which was performed using gentle suction via a small pipette (~3-4 μm in diameter). This procedure is widely used to gain access to the different cochlear cell types, including the OHCs (**Marcotti & Kros, 1999**) and it does not affect their integrity, since OHCs retained the ability to generate Ca^2+^ transients (**Supplementary Figure 2A**) and sensitivity to extracellular ATP (**Supplementary Figure 2B**). After the removal of the Deiters’ cells, we elicited Ca^2+^ waves by photo-damaging a small area in the GER at room temperature. This treatment was used as a proxy for spontaneous Ca^2+^ waves (**Lahne and Gale, 2010**) because it mimics the influence on the OHCs (**Figure 5A,B, Movie 6**). We found that Ca^2+^ elevation in OHCs associated with Ca^2+^ waves in the GER was almost completely abolished when Deiters’ cells were removed (**Figure 5C-E:** *P*<0.001 compared to when Deiters’ cells were present; post-test from one-Way ANOVA) or in the absence of Ca_V_1.3 Ca^2+^ channels (**Figure 5C-E**). We conclude that Deiters’ cells are essential intermediaries for coupling activity from the GER to OHCs.

**Figure 4.**
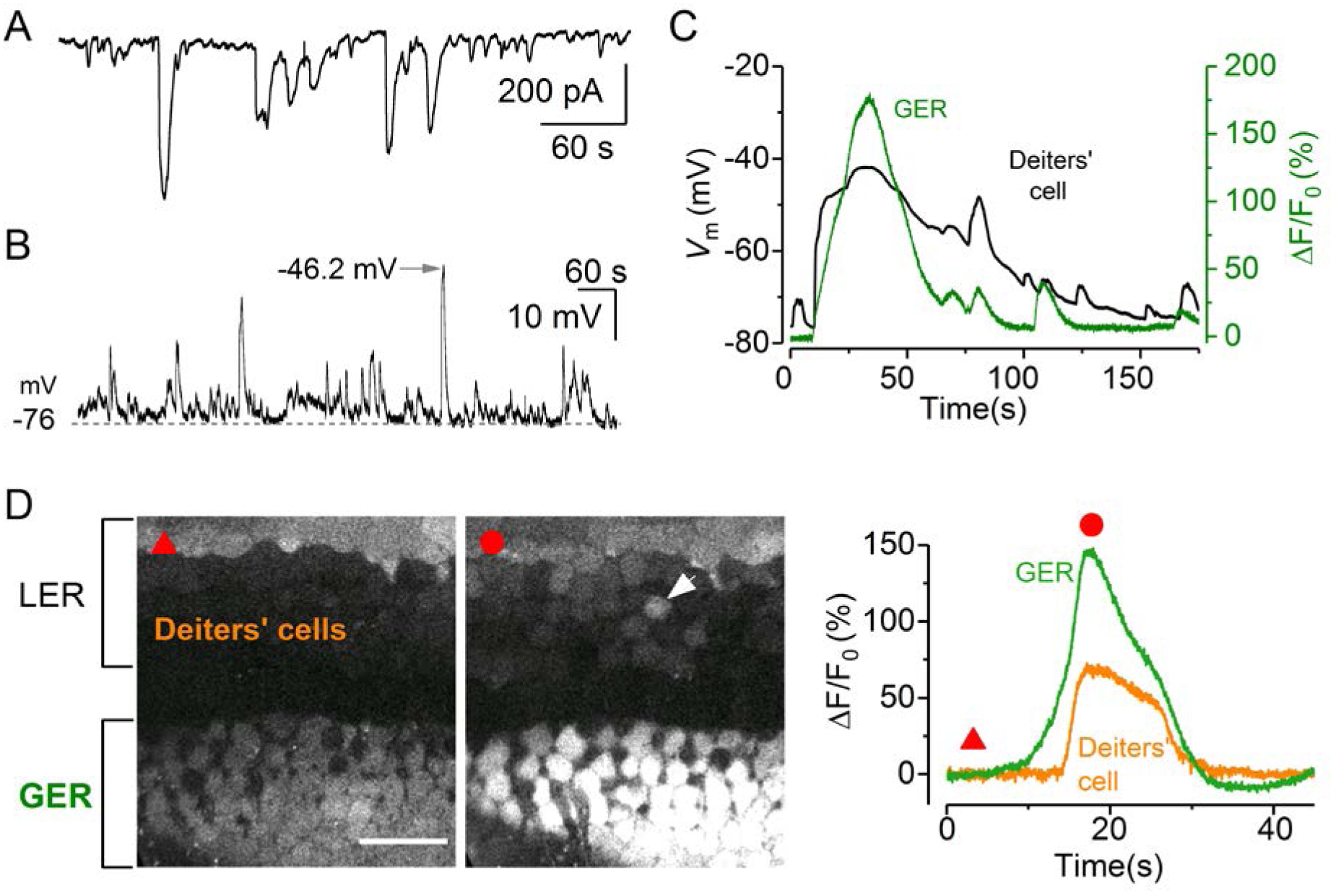
Calcium signalling from the GER travels to Deiters’ cells in the LER. (**A** and **B**) Spontaneous activity recorded using the patch clamp technique from Deiters’ cells using whole-cell voltage clamp (**A**) and current clamp (**B**) in a P1 mouse cochlea. Recordings are at RT. (**C**) Simultaneous recording of whole-cell voltage responses in a Deiters’ cell in the LER (black trace) and a spontaneous Ca^2+^ wave from the GER (ΔF/F0) of a *Ca_V_1.3^-/-^* P1 mouse (for wild-type mouse cochlea see **Supplementary Figure 1**). (**D**) Representative ΔF/F0 traces (right traces) from the GER and one Deiters’ cell (arrowhead in the middle panel) of a P1 *Ca_V_1.3^-/-^* mouse. Note that the response in the GER occurs earlier than that in the Deiters’ cell (left), even though the rapid onset of the voltage change in the Deiters’ cell gives the impression of an overlapping event (**C**). Traces are computed as pixel averages of regions of interest.

**Figure 5.**
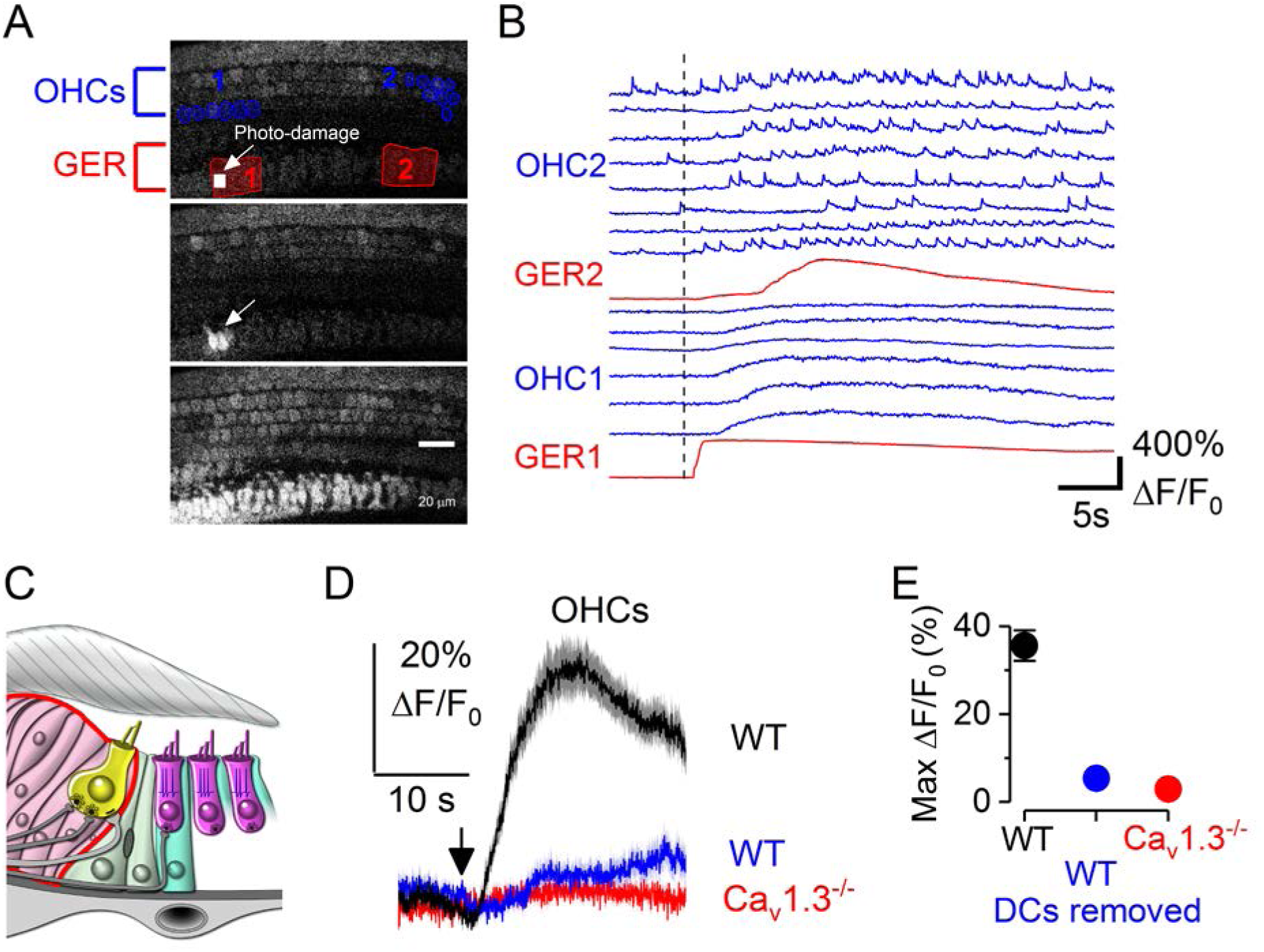
Deiters’ cells are required for the modulation of OHC APs by Ca^2+^-signalling from the GER. (**A**) Three representative images showing the small photo-damage region in the GER (top panel, small white rectangles - arrow), the initiation of the Ca^2+^ wave from the damaged non-sensory cells (middle panel: arrow) and the full extent of the Ca^2+^ wave (bottom panel). Note that during the Ca^2+^ wave, the OHCs in the LER also become bright. The red areas or regions of interest in the top panel are the region used to measure Ca^2+^ signals at the point where the Ca^2+^ wave was generated (GER1) and its propagation along the epithelium (GER2). The blue ROIs are those used to measure OHC Ca^2+^ signalling. Recording were performed at RT from wild-type P2 mouse. The photo-damage area was 55 μm^2^ in size (18 × 23 pixels) and typically covered the apical surface of ~1-2 non-sensory cells in the GER. (**B**) Representative ΔF/F0 traces from the GER (red traces) and OHCs (blue traces). Note that the 6 OHCs traces named “OHC1” are those closer to GER1 (photo-damage region: see top panel **A**), while the 8 OHC traces named “OHC2” are those near GER2 (far away from the photo-damage region: see top panel **A**). Recordings are at RT. (**C**) Diagram showing the immature organ of Corti without the Deiters’ cells. Note that for these experiments two (as shown in the diagram) or all three rows of Deiters’ cells (usually spanning the distance of 5-10 OHCs) were removed prior to performing the Ca^2+^ imaging experiments. (**D** and **E**) Average (**D**) and maximum (**E**) Ca^2+^ responses from apical OHCs (P2-P3) induced by photo-damage of non-sensory cells in the GER. Black trace and symbol (wild-type cochlea with Deiters’ cells) are average of 85 OHCs from 9 recordings; blue trace and symbol (wild-type cochlea in which one or no Deiters’ cells were present: for simplicity one cell is shown in the image in panel **C**) are average of 72 OHCs from 9 recordings; red trace and symbol (*Ca_V_1.3^-/-^* cochlea with Deiters’ cells but with electrically silent OHCs) are average of 175 OHCs from 10 recordings.

### ATP triggers OHC electrical activity in the developing cochlea

To determine the molecular mechanism linking activity in the Deiters’ cells with OHC synchronization, we pharmacologically probed the basolateral membrane of OHCs deprived of their surrounding Deiters’ cells (**Figure 6A**). Deiters’ cells release ATP via connexin hemichannels (**Zhao et al., 2005**) and immature OHCs exhibit depolarizing, ATP-gated currents (**Glowatzki et al., 1997**). In the absence of Deiters’ cells, we found that local perfusion of 10 μM ATP onto the basolateral membrane of OHCs triggered large Ca^2+^ responses (**Figure 6B**). The OHC response to ATP was abolished in *Ca_V_1.3^-/-^* mice even when used at 100 μM (**Supplementary Figure 3, Movie 7**). Under whole-cell patch clamp, 10 μM and 100 μM ATP caused OHCs to depolarize by 15.8 ± 2.9 mV (steady-state, *n* = 10, P1-P2) (**Figure 6C; Supplementary Figure 3**). Extracellular ATP can act on ionotropic (P2X) and metabotropic (P2Y) purinergic receptors, both of which are present in cochlear hair cells (**Housley et al., 2006**). We found that Ca^2+^ signals from OHCs were either abolished or greatly reduced when ATP was applied together with the purinergic receptor antagonists suramin (200 μM: **Figure 6D,I**) and PPADS (**Figure 6I**). Under the whole-cell patch clamp configuration, suramin reduced the ATP responses by 89.8 ± 6.5 % (*n* = 4, P1) (**Figure 6C**). The absence of ATP-induced intracellular Ca^2+^ signals in wild-type OHCs bathed in a Ca^2+^-free solution (**Figure 6E,I**) and in *Ca_V_1.3^-/-^* mice (**Figure 6I, Supplementary Figure 3, Movie 7**) indicates that P2Y receptors, which mobilize Ca^2+^ from intracellular stores, are unlikely to be involved in mediating OHC responses to ATP at this developmental stage (**King and Townsend-Nicholson, 2003; Egan & Khakh, 2004**). Consistent with this hypothesis, the application of the phospholipase C inhibitor U73122, which prevents the IP_3_-mediated Ca^2+^ release from intracellular Ca^2+^ stores linked to the activation of P2Y receptors (**Bleasdale and Fisher, 1993; Lahne and Gale, 2010**), did not inhibit ATP induced Ca^2+^ signals in OHCs (**Figure 6F,I**). The local application onto Deiters’ cells of 1 μM UTP, which is a selective agonist for P2Y receptors and is involved in intercellular Ca^2+^ signals in the non-sensory cells including the Deiters’ cells (**Piazza et al., 2007**), increases their intracellular Ca^2+^ signals, followed by increased Ca^2+^ transients in nearby OHCs (**Figure 6G, Movie 8**). When the Deiters’ cells were removed, OHCs were not affected by UTP (**Figure 6H**: normalized maximal response: DCs intact: 1.00 ± 0.23, *n* = 4 recordings, 3 cochleae, 3 mice; DCs removed: 0.20 ± 0.04, *n* = 5 recordings, 3 cochleae, 3 mice; *P*<0.01, Mann-Whitnhey U test), providing further evidence that ATP-induced Ca^2+^ signals from these non-sensory cells directly modulates OHC activity. Altogether these data indicate that ATP, acting through P2Y receptors in Deiters’ cells and P2X receptors in OHCs, can coordinate the activity of nearby OHCs.

**Figure 6.**
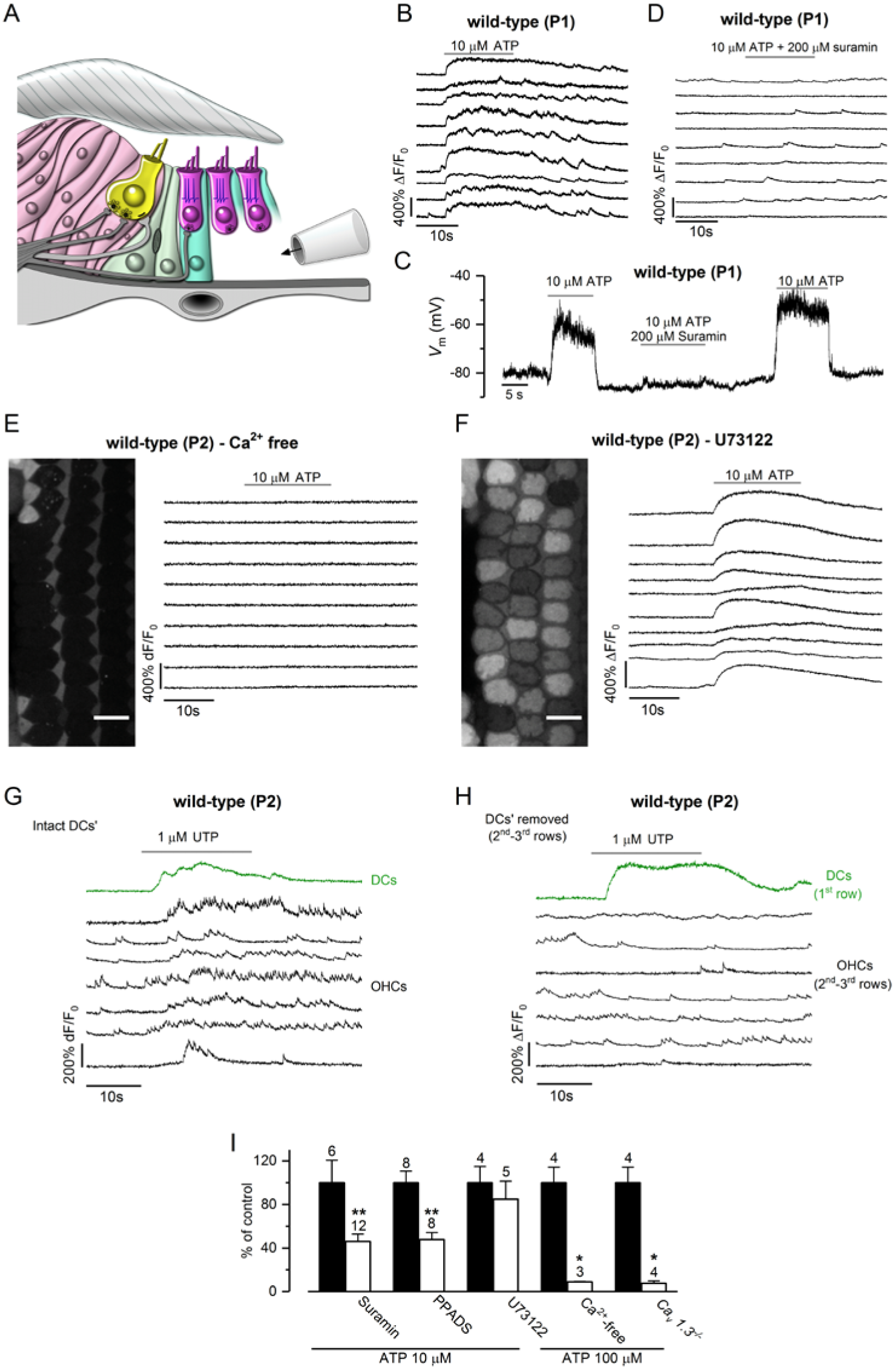
ATP-induced modulation of the OHC firing activity is mediated by ionotropic purinergic receptors. (**A**) Diagram showing a portion of the immature organ of Corti highlighting the experimental approach used to locally perfuse channel blockers and ATP directly to the basolateral membrane of OHCs *in situ*. (**B**) Representative ΔF/F0 traces from 9 apical OHCs of a P1 wild-type mouse during the application of 10 μM ATP. Data analysis as in **Figure 1C**. (**C**) Voltage responses in whole-cell current clamp from a P1 OHC of a wild-type mouse during the extracellular application of 10 μM ATP alone or together with 200 μM of the P2X receptor blocker suramin. Suramin largely reduce the ATP-induced OHC depolarization. (**D**) Representative ΔF/F0 traces from 9 apical OHCs of a P1 wild-type mouse during the application of 10 μM ATP + 200 μM suramin. Note that suramin prevents the occurrence of the large Ca^2+^ signals in OHCs. Data analysis as in **Figure 1*C***. (**E** and **F**). Representative ΔF/F0 traces from 10 apical OHCs of P1 wild-type mice (selected from the images in the left panel) during the application of 10 μM ATP in the absence of Ca^2+^ in the extracellular solution (**E**) or in the presence of the metabotropic P2Y receptor blocker U73122 (10 μM, **F**). Note that large OHC depolarizations obtained with ATP cause sustained Ca^2+^ signals. (**G** and **H**) The P2Y agonist UTP (1 μM) causes increased Ca^2+^ signals in the Deiters’ cells, which directly elevate the firing activity of OHCs (**G**). Note that in the absence of Deiters’ cells (**H**) OHC did not respond to UTP when deprived of Deiters’ cells. (**I**) Histogram showing maximum ΔF/F0 Ca^2+^ responses in P1-P2 OHCs to the extracellular application of ATP (10 μM or 100 μM) alone (black columns) and together with the purinergic receptor blockers suramin (200 μM) PPADS (100 μM) and U73122 (10 μM), Ca^2+^-free extracellular solution and in *Ca_v_1.3^-/-^* mice. Experiments were performed in the absence of the Deiters’ cells. Responses in each condition are normalised to control experiments, carried out under the same imaging and dye-loading conditions. Number of recordings shown above the columns. *p<0.05, **p<0.01, Mann-Whitney U test.

### The frequency of large Ca^2+^ waves is reduced in *Cx30^-/-^ mice*

In the sensory epithelium of the mammalian cochlea, gap junctions are formed by connexin 26 (Cx26) and Cx30 (**Lautermann et al., 1998**). To test the role of gap junctions in the spread of spontaneous Ca^2+^ activity, we used *Cx30^-/-^* mice (**Teubner et al., 2003**) in which the mRNA and protein expression of Cx30 are abolished and those of Cx26 are reduced to only ~10% of normal levels during pre-hearing stages (**Boulay et al., 2013**). Despite the loss of connexins, spontaneous and rapid Ca^2+^-dependent APs in developing OHCs were still recorded (**Figure 7A**). These Ca^2+^ spikes occurred in both apical and basal OHCs as shown in wild-type mice (**Figure 1**) and the number of active OHCs decreased with age (**Figure 7B**). Thus the intrinsic Ca^2+^ firing activity in developing OHCs was unaffected by the loss of connexins in the non-sensory cells. In agreement with previous findings (**Rodriguez et al., 2012**), the average frequency of Ca^2+^ waves in the GER of *Cx30^-/-^* mice (1.3 ± 0.1 events/min, *n* = 36 recordings, 11 cochleae, 11 mice, P1-P2) was significantly reduced compared to that of wild-type mice (2.09 ± 0.21 events/min, *n* = 32 recordings, 7 cochleae, 6 mice, P1-P2, *P*=0.003, Mann-Whitney U-test). Furthermore, the frequency of the larger Ca^2+^ events (>75μm: see **Figure 3**) required for the synchronization of several OHCs was reduced ~5 fold in *Cx30^-/-^* (0.10 ± 0.03 events/min, **Figure 7C**) compared to wild-type (0.53 ± 0.11 events/min, **Figure 3**, *P*<0.0001). The few remaining large Ca^2+^ waves in *Cx30^-/-^* mice were still able to synchronize the electrical activity of OHCs (**Figure 7C**). Overall, these findings show that the Ca^2+^ signalling from non-sensory cells, although not required for generating spontaneous APs in OHCs, is crucial to synchronize their activity.

**Figure 7.**
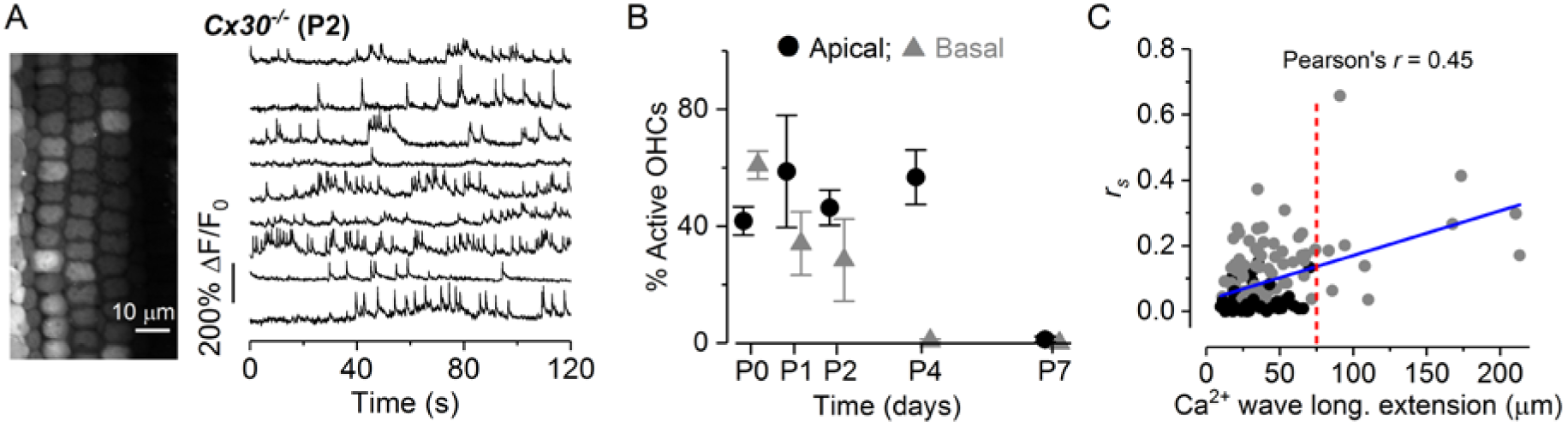
Calcium waves from the GER modulate OHCs spontaneous Ca^2+^-signalling. (**A**) Representative ΔF/F0 traces from 9 apical OHCs of a P2 Cx30^-/-^ mouse (selected from the images in the left panel) recorded at 31°C. Data analysis as in **Figure 1C**. (**B**) Percentage of apical and basal OHCs showing spontaneous Ca^2+^ spikes as a function of postnatal age. Number of total OHCs and recordings from left to right were: apical cochlea 404 and 7; 156 and 3; 368 and 7; 454 and 8; 365 and 8; basal cochlea 441 and 8; 375 and 6; 328 and 7; 487 and 9; 346 and 6. (**C**) Average Spearman’s rank correlation coefficient (*r*_s_) between the OHC activity as a function of the longitudinal extension of spontaneous Ca^2+^ wave in the GER from the apical coil of P1-P2 mouse cochleae (for additional details see **Figure 3D**). Solid line represents a linear fit to the data. The slope was significantly different from zero (*P*<0.0001)

### The biophysical characteristics of OHCs from *Cx30*^-/-^ mice develop normally

In OHCs the onset of maturation occurs at around P7-P8 when they begin to express a negatively-activated K^+^ current *I*_K,n_ and acquire electromotile activity (**Marcotti and Kros, 1999; Abe et al., 2007**). The expression of the motor protein prestin (**Zheng et al., 2000; Liberman et al., 2002**), which drives the somatic motility of OHCs was normal between wild-type and *Cx30^-/-^* mice (see **Figure 10**). The maturation of OHCs is also associated with an increase in cell membrane capacitance (**Marcotti and Kros, 1999**), which was observed in both genotypes (**Figure 8C**: *P* = 0.4350). The total outward K^+^ current (*I*_K_) and the isolated *I*_K,n_ recorded from OHCs of *Cx30^-/-^* mice (P10-P12) were similar in size to that of wild-type cells (*I*_K_: *P* = 0.8445; *I*_K,n_: *P* = 0.2238, **Figure 8C**). Mature OHCs are the primary target of the inhibitory olivocochlear efferent fibres that release the neurotransmitter acetylcholine (ACh) (**Simmons et al., 1996**). Efferent inhibition of OHCs by ACh is achieved by Ca^2+^ influx through α9α10-nAChRs activating a hyperpolarizing SK2 current (**Oliver et al., 2000; Marcotti et al., 2004; Lioudyno et al., 2004; Katz et al 2004**). Mouse OHCs first become highly sensitive to ACh from around the end of the first postnatal week (**Marcotti et al., 2004; katz et al., 2004**), which coincides with their onset of functional maturation (**Marcotti & Kros, 1999**). In the presence of ACh, depolarizing and hyperpolarizing voltage steps from a holding potential of –84 mV elicited an instantaneous current in wild-type OHCs. This ACh-activated instantaneous current is mainly carried by SK2 channels but also by nAChRs since it is blocked by apamin and strychnine, respectively (**Marcotti et al., 2004**). The ACh-activated current was present in OHCs from wild-type (**Figure 8D**) and *Cx30^-/-^* mice (**Figure 8*E***). The sensitivity of OHCs to ACh was quantified by measuring the steady-state slope conductance at –84 mV of the ACh-sensitive current (*g*_ACh_), which was obtained by subtracting the control currents from the currents in the presence of 100 μM ACh (**Figure 8D,E: see also Marcotti et al.,2004**). *g*_ACh_ was similar between wild-type (8.4 ± 1.7 nS, *n* = 4, P12) and *Cx30^-/-^* (8.4 ± 1.4 nS, *n* = 6, P10-P12; P*=*0.9843). We further confirmed that the ACh-induced currents in *Cx30^-/-^* OHCs were carried by the nAChRs and SK2 channels since they were blocked by strychnine (at –90 mV: **Figure 8F**) and a Ca^2+^ free solution (at –40 mV: **Figure 8G**), respectively, as previously shown in hair cells (**Glowatzki and Fuchs, 2000; Oliver et al., 2000; Marcotti et al., 2004**).

**Figure 8.**
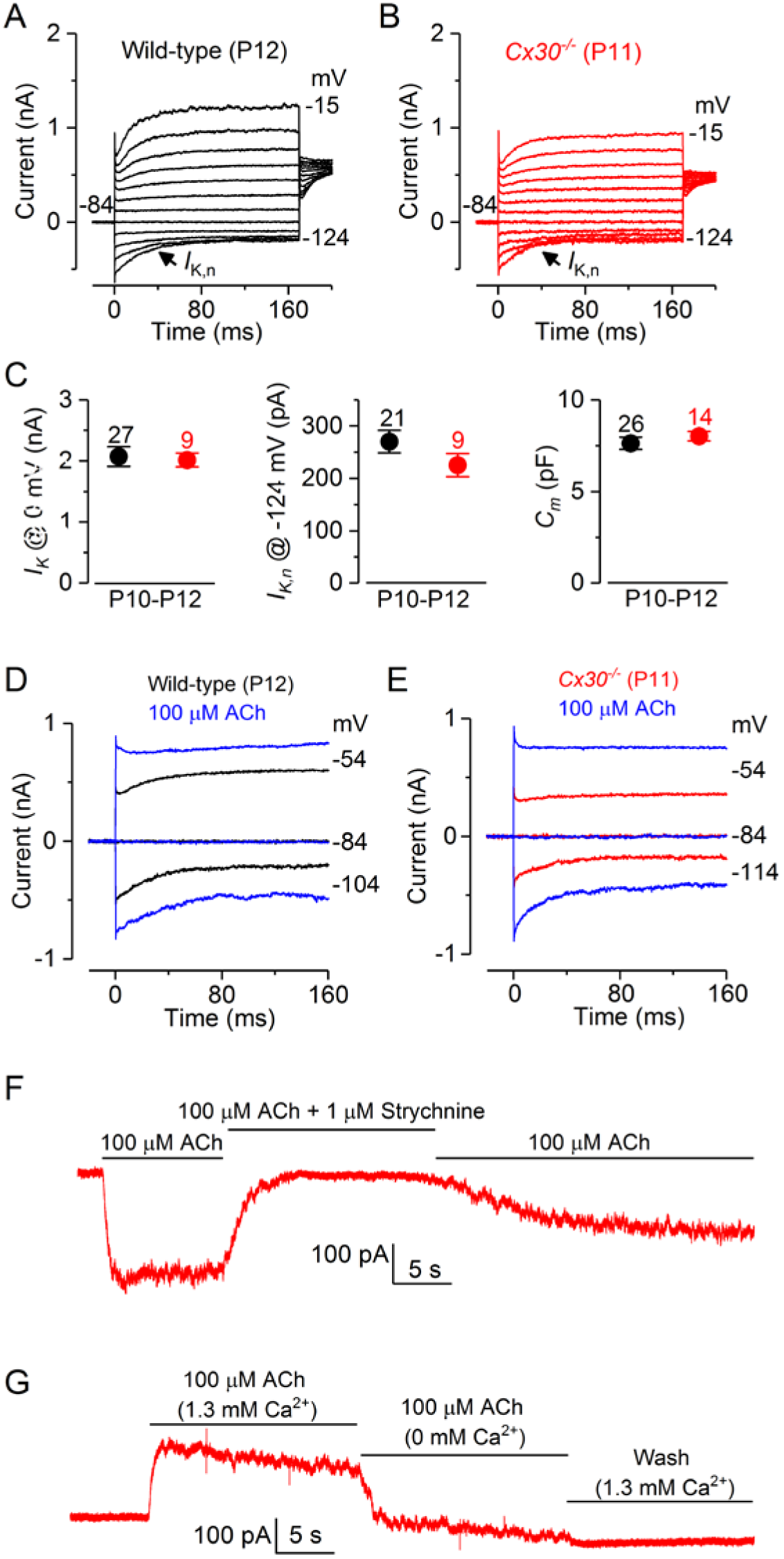
OHCs from mature Cx30^-/-^ mice develop normal biophysical properties. (**A, B**) Current responses in wild-type (**A**) *Cx30^-/-^* (**B**) apical coil OHCs after their onset of maturity, which occurs at P7-P8. Outward current was elicited by using depolarizing and hyperpolarizing voltage steps (10 mV increments) from –84 mV to the various test potentials shown by some of the traces. (**C**) Average amplitude to the total outward current (*I*_K_: left), the isolated *I*_K,n_ (middle) and the membrane capacitance (*C*_m_: right) of P10-P12 OHCs. (**D,E**) Membrane currents recorded from OHCs in wild-type (**D**, P12) and *Cx30^-/-^* (**E**, P11) mice before and during superfusion of 100 μM ACh. (**F**) In *Cx30^-/-^* OHCs, the inward current elicited in 100 μM extracellular ACh at –90 mV was reversible blocked by 1 μM strychnine, indicating the direct involvement of α9α10nAChRs. (**G**) At –40 mV, the outward current in *Cx30^-/-^* OHCs was prevented by an absence of Ca^2+^ in the extracellular solution, indicating the presence of SK2 channels.

### OHC ribbon synapses and afferent fibres are reduced in *Cx30^-/-^* mice

The above results demonstrate that pre-hearing *Cx30^-/-^* mice, in which OHCs retain their intrinsic AP activity but have a reduced number and more confined Ca^2+^-wave in the GER, were able to acquire functionally mature OHCs from a biophysical point of view. During the same time window (~P0-P12), immature IHCs from *Cx30^-/-^* mice have been shown to be normal (**Johnson et al., 2017**), indicating that Ca^2+^-waves in the non-sensory cells do not interfere with the normal pre-hearing development of hair cells. Indeed, it has been suggested that the modulation of AP activity in IHCs by the Ca^2+^-waves could be used to refine the afferent auditory pathway (**Tritsch et al., 2007**), although direct proof for this is still missing. Therefore, we made use of the fact that hair cells from *Cx30^-/-^* mice are normal during pre-hearing stages, to investigate whether the Ca^2+^-signalling in the GER contributes to the refinement of OHC afferent innervation.

The OHC afferent ribbon synapses from wild-type and *Cx30^-/-^* mice were investigated before (P4) and after (P10) their onset of functional maturation at ~P8 (**Simmons, 1994**). At P4 both wild-type and *Cx30^-/-^* OHCs showed a similar number of ribbons (*P* = 0.49: **Figure 9A,B,E:** CtBP2 puncta in red; Myo7a, blue, was used as the hair cell marker). In mature OHCs (P10) the number of ribbons in *Cx30^-/-^* OHCs was about half to that of wild-type OHCs (*P* < 0.0001: **Figure 9C,D,E**). As a comparison, we also looked at IHCs and found a similar number of ribbons between the two genotypes at both P4 (*P* = 0.30: **Figure 9F,G,J**) and P10 (*P*=0.43: **Figure 9H,I,J**), further supporting the evidence that at this age immature IHCs are unaffected by the absence of Ca^2+^ waves (**Johnson et al., 2017**). We then looked at whether the reduction in ribbon synapses was also associated with anomalies in the afferent fibres innervating mature OHCs. Prestin immunoreactivity was used as the OHC marker (**Figure 10A,B**). At P11 in the apical cochlear region, afferent fibres form spiral ganglion neurons form outer spiral fibres that terminate on the OHCs after long spiral courses (**Simmons and Liberman, 1988**). These fibres show peripherin immunoreactivity (**Hafidi, 1998**) and course radially from the spiral ganglion to the organ of Corti, cross along the floor of the tunnel of Corti and spiral in a basal direction before giving rise to punctate endings on OHCs (**Figure 10C**). In the apical region, there were 14.0 ± 2.0 (mean ± S.D., *n* = 3 mice) tunnel-crossing outer spiral fibres per 100 μm along the length of the organ of Corti. Compared to wild-type, *Cx30^-/-^* mice had fewer peripherin-labelled outer spiral fibres (**Figure 10D**), which was matched by a reduction in labelled fibres crossing the tunnel of Corti (7.0 ± 1.0 fibres per 100 μm, *n* = 3 animals) compared to wild-type mice (*P* <0.01). This result is in agreement with the significantly lower number of ribbon synapses in OHCs from *Cx30^-/-^* mice (**Figure 9E**).

**Figure 9.**
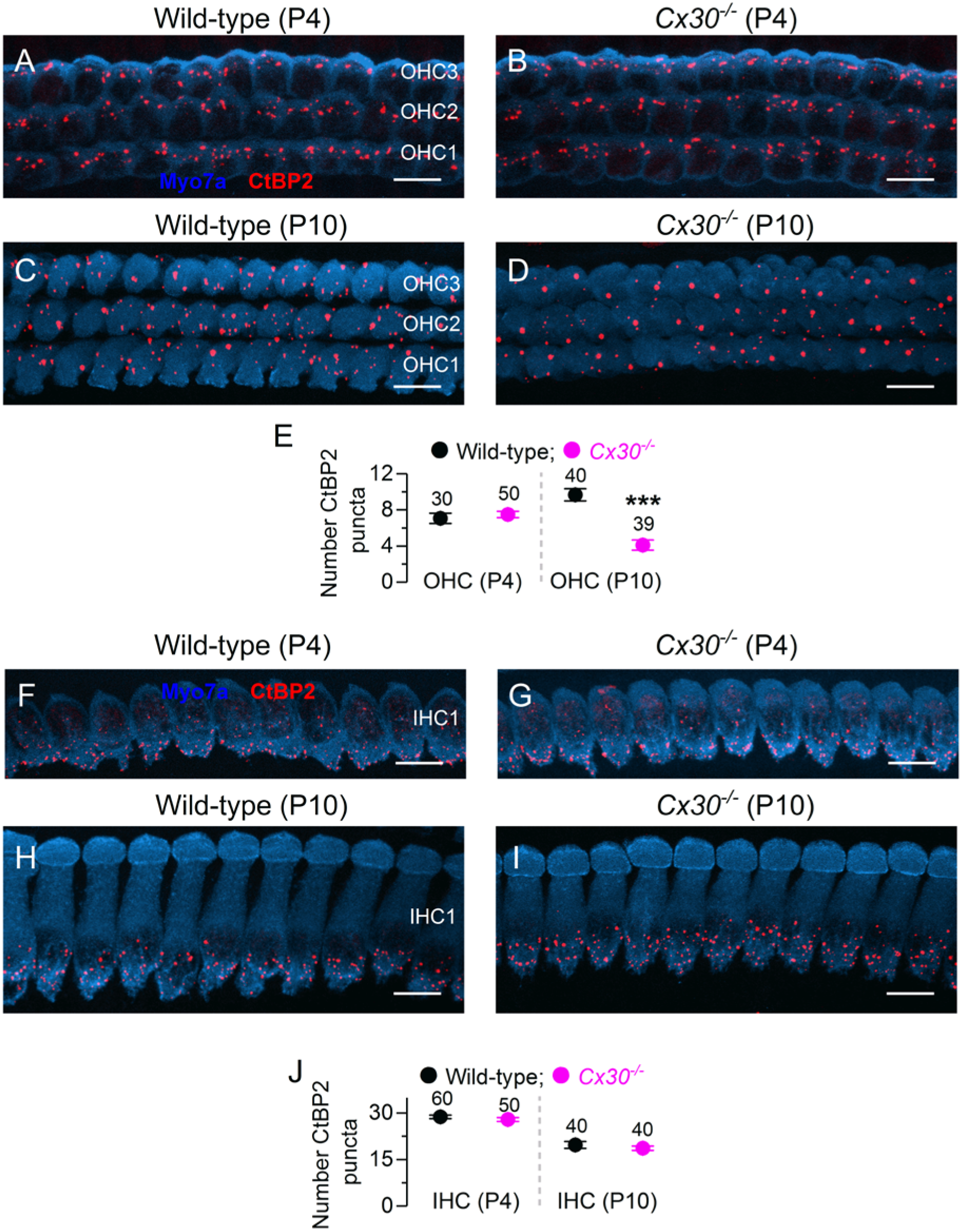
Ribbon synapses are reduced in *Cx30^-/-^*OHCs. (**A-D**) Maximum intensity projections of confocal z-stack images that were taken from apical coil OHCs before (P4) and after (P10) their onset of functional maturation at P8 in wild-type (**A** and **C**) and *Cx30^-/-^* (**B** and **D**) mice. Immunostaining for ribbon synapses (CtBP2) is shown in red; Myo7a (blue) was used as the hair cell marker. (**E**) Number of ribbon (CtBP2 puncta) in wild-type and *Cx30^-/-^* OHCs at P4 and P10. (**F-J**) Maximum intensity projections of confocal z-stack images (**F-I**) and ribbon count (**J**) from IHCs. Experimental conditions as in panels **A-E**. Number of OHCs and IHCs analysed is shown above each average data point; 4 mice were used for each experimental condition. *** indicates *P* < 0.0001. Scale bars 10 μm. Note that the size or ribbon (CtBP2 puncta) in OHCs (**A-D**) are normally larger than that seen in IHCs (**F-I**), which is likely to reflect the multiple synaptic ribbons at OHC active zones (Shnerson *et al.* 1982).

**Figure 10.**
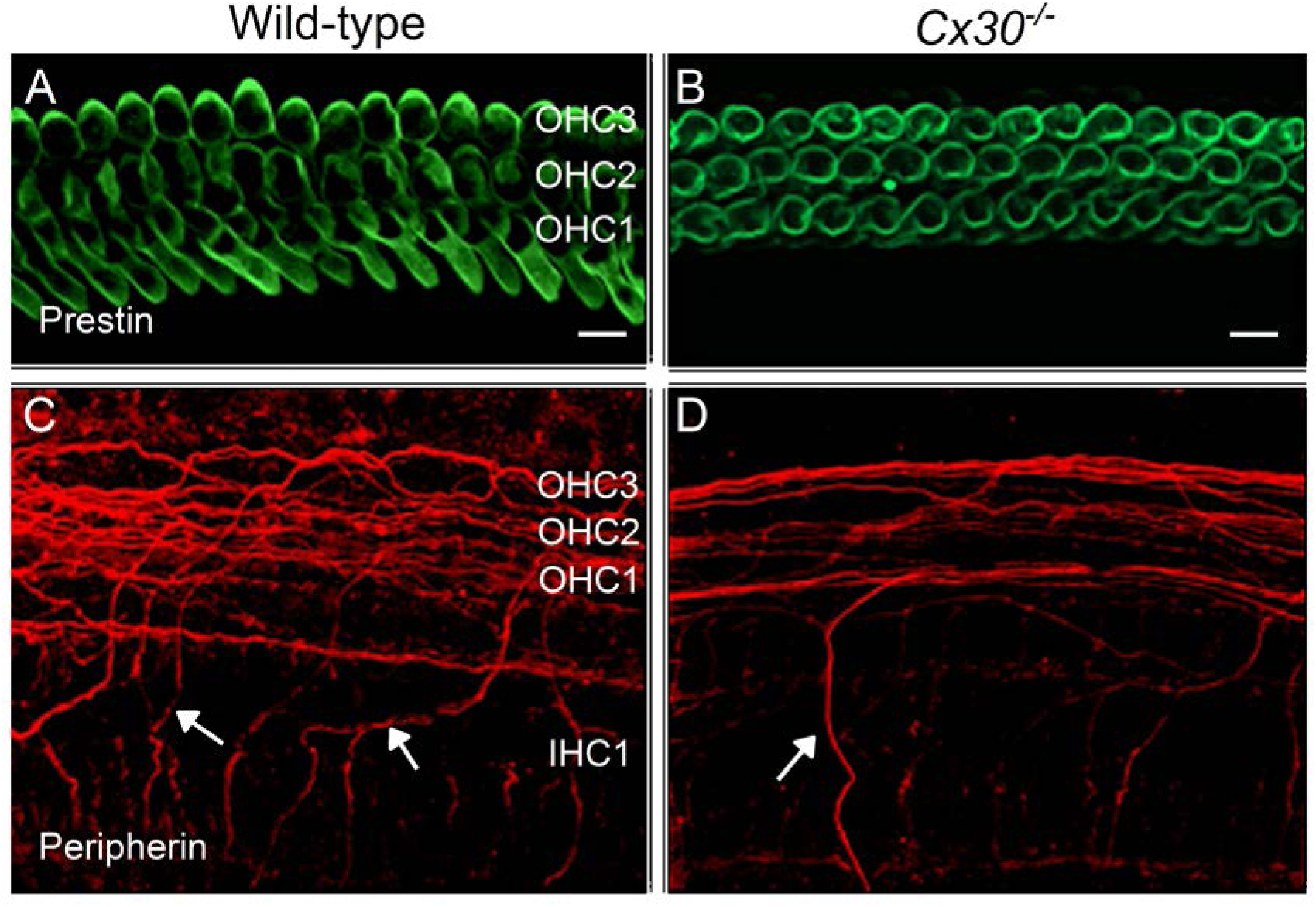
Afferent fibres are reduced in *Cx30^-/-^* mice. Maximum intensity projections of confocal z-stacks taken from the apical cochlear region of wild-type (left column) and *Cx30^-/-^* (right column) mice at P11 using antibodies against prestin (green) and peripherin (red). Each panel represents a different mouse. (**A** and **B**) Prestin labelling was similar between the different mouse strains and as such was used as an OHC marker. Scale bars 10 μm. (**C** and **D**) Immunostaining for peripherin (red) highlights outer spiral fibres (arrows) of type II spiral ganglion neurons in the wild-type mouse cochlea (**D**). These outer spiral fibres cross below IHCs and spiral below OHCs towards the cochlear base. In *Cx30^-/-^* (**D**) mice there are fewer peripherin-labelled outer spiral fibres than in wild-type.

## Discussion

We have identified distinct, coordinated Ca^2+^-dependent mechanisms that influence the refinement of OHC innervation. Our evidence shows that the morphological maturation of the afferent synapses and innervation of OHCs requires spontaneous ATP-induced Ca^2+^ waves in the non-sensory cells of the GER, which synchronise the action potential (AP) activity between several OHCs. The immature cochlea uses similar Ca^2+^-dependent signalling to drive the maturation of IHCs (**Johnson et al., 2013**) but does not appear to affect their innervation pattern. Moreover this Ca^2+^ activity, which different from that influencing IHC APs (**Wang et al., 2015**) is mediated by ATP-induced activation of P2X receptors, has distinct functional consequences for OHC maturation and is separated by developmental timing, with OHCs preceding IHCs (**Johnson et al., 2011; 2017**). The data suggest that several distinct patterns of spontaneous, experience-independent Ca^2+^ activity across the auditory sensory epithelium orchestrate the differential maturation of OHCs (afferent innervation) and IHCs (sensory cells: **Johnson et al., 2013**) to shape the final stages of auditory organ development.

### OHC activity is synchronised by ATP-induced Ca^2+^ signalling in non-sensory cells

We show that spontaneous intercellular Ca^2+^ signalling activity originating in the non-sensory cells of the greater epithelial ridge (GER) synchronizes Ca^2+^ AP activity between nearby OHCs via release of ATP from the Deiters’ cells. This ATP acts directly via P2X receptors on the OHCs. The same activity in the GER synchronizes APs in IHCs (**Tritsch et al., 2007; Johnson et al., 2011; Wang et al., 2015**) but using a different mechanism. In IHCs, ATP indirectly increases the firing activity of IHCs by acting on purinergic autoreceptors expressed in the non-sensory cells surrounding the IHCs, which leads to the opening of TMEM16A Ca^2+^-activated Cl^-^ channels and the efflux of K^+^ in the intercellular space (**Wang et al., 2015**). The expression of these TMEM16A channels seems to follow closely the development of IHCs and they are absent in the LER (**Wang et al., 2015**). Although P2X_2_ are the most abundant purinergic receptors in the cochlea, they are mainly expressed in hair cells from the second postnatal week onward throughout adult stages (**Järlebark et al., 2000**), so they are unlikely to mediate the ATP-induced signalling in developing OHCs. P2X_4_ receptors have been suggested to be present in the developing cochlea based on pharmacological assays, but these findings have not been confirmed with expression studies (**Lamne and Gale, 2010**). Of the other known P2X receptors, only P2X_3_ (**Huang et al., 2006**) and P2X_7_ (**Nikolic et al., 2003**) have been shown to be transiently expressed during early stages of development. Indeed, the expression time course of P2X_3_ receptors seems to match our Ca^2+^-imaging experiments (**Figure 1**), since a previous study has shown that by P3 they are still present in apical, but no longer in basal OHCs (**Huang et al., 2006**). P2X_3_ receptors have also been implicated in early development in the peripheral and central nervous system (**Kidd et al., 1998**).

Although ATP-induced Ca^2+^ signalling from non-sensory cells is crucial for promoting the maturation of IHCs after the onset of hearing (**Johnson et al., 2017**), our data suggests that it does not contribute directly to OHC maturation. However, the time course of maturation of OHCs and IHCs is very different (**Corns et al., 2014**). During the time over which OHCs become sensory competent, IHCs are still immature and driving the functional refinement of the auditory pathway, which mainly occurs during the first week of postnatal development (**Snyder and Leake, 1997; Kim and Kandler, 2003; Zhang-Hooks et al., 2016**). IHCs begin to mature towards the end of the second postnatal week (at the onset of hearing), indicating that Ca^2+^ waves from the non-sensory cells are likely to drive different signals to the hair cells during early and later stages of pre-hearing development. There are thus temporally distinct critical windows for the influence of spontaneous Ca^2+^ activity in non-sensory cells on IHCs and OHCs.

### OHC afferent innervation is shaped by ATP-induced intercellular Ca^2+^ signalling in non-sensory cells

In the cochlea, the onset of OHC function is associated with Type II spiral ganglion afferent terminals forming extensive arborisations with several OHCs (**Perkins and Morest, 1975; Echteler, 1992**). Unlike Type I afferent fibres contacting IHCs, Type II afferent fibres seem to only respond to the loudest sounds (**Robertson, 1984; Browm, 1994**), which has led to the assumption that they represent the cochlear nociceptors (**Weisz et al., 2009; Liu et al., 2015**). Our imaging experiments show that large spontaneous Ca^2+^ waves originating in the GER (~200 μm in longitudinal extension) are able to synchronize bursting activity between OHCs. Considering that the average spiral processes of type II fibres span 215 μm (**Weisz et al., 2012; Martinez-Monedero et al., 2016**) and that they contact more than a dozen OHCs (**Perkins and Morest, 1975**), these Ca^2+^ waves should be sufficient to activate most of the presynaptic, immature OHCs that form synapses with each developing afferent fibre. Since OHCs provide an infrequent and weak synaptic input to Type II afferent fibres, their suprathreshold excitation would require the summation of the input coming from all OHCs contacting each fibre (**Weisz et al., 2009;2012**), which in the developing cochlea could be provided by the Ca^2+^ waves originating in the GER. The synchronized activity among nearby OHCs would lead to periodic stimulation of the Type II afferent fibres and leading to activity-dependent refinement of synaptic connections as also seen in the visual system (**Katz & Shatz,1996; Spitzer, 2006**). Indeed, we found that an absence of connexins in non-sensory cells, which reduces the frequency and spatial extent of the Ca^2+^ waves and as such OHC synchronization (*Cx30^-/-^* mice**: Figure 7**), lead to a reduced number of ribbon synapses and Type II afferent fibres. A similar phenotype in the Type II afferent innervation was also seen in mice lacking Deiters’ cells (**Mellado Lagarde et al., 2013**), corroborating our finding that these non-sensory cells are crucial for the transfer of information from the GER to the OHCs.

In summary, we propose that in the immature mammalian cochlea the refinement of the OHC afferent innervation pattern requires synchronized activity between neighbouring OHCs, which is provided via Deiters’ cells from large Ca^2+^ waves originating in the GER. Overall our results reveal extraordinary physiological regulation of spontaneous Ca^2+^ signalling in the developing cochlea over discrete and separate time periods, to ensure the correct functional differentiation of neuronal and sensory cells in the maturing auditory system.

## Materials and Methods

### Ethics Statement

The majority of the animal studies were performed in the UK and licensed by the Home Office under the Animals (Scientific Procedures) Act 1986 and were approved by the University of Sheffield Ethical Review Committee. Some experiments were performed in the USA and the animal work was licensed by the Baylor University IACUC (Institutional Animal Care and Use Committee) as established by U.S. Public Health Service.

### Tissue preparation

Apical- and basal-coil OHCs from wild-type mice or transgenic mice of either sex were studied in acutely dissected organs of Corti from postnatal day 0 (P0) to P13, where the day of birth is P0. Transgenic mice include *Cx30^-/-^* (MGI:2447863) (**Teubner et al., 2003**) and *Ca_V_1.3^-/-^* mice (**Platzer et al., 2000**). The genotyping protocols for these transgenic mice were performed as previously described (**Teubner et al., 2003; Platzer et al., 2000**). Mice were killed by cervical dislocation and the organ of Corti dissected in extracellular solution composed of (in mM): 135 NaCl, 5.8 KCl, 1.3 CaCl_2_, 0.9 MgCl_2_, 0.7 NaH_2_PO_4_, 5.6 D-glucose, 10 Hepes-NaOH. Sodium pyruvate (2 mM), amino acids and vitamins were added from concentrates (Thermo Fisher Scientific, UK). The pH was adjusted to 7.5 (osmolality ~308 mmol kg^-1^). The dissected organ of Corti was transferred to a microscope chamber, immobilized using a nylon mesh fixed to a stainless steel ring and viewed using an upright microscope (Olympus BX51, Japan; Leica, DMLFS, Germany; Bergamo II System B232, Thorlabs Inc.). Hair cells were observed with Nomarski differential interface contrast optics (x63 water immersion objective) or Dodt gradient contrast (DGC) optics (x60 water immersion objective) and either x10 or x15 eyepieces.

### Single-cell electrophysiology

Membrane currents and voltage responses were investigated either at room temperature (20-24 °C) or at body temperature (33-37 °C), using Optopatch (Cairn Research Ltd, UK) or Axopatch 200B (Molecular Devices, USA) amplifiers. Patch pipettes, with resistances of 2-3 MΩ, were pulled from soda glass capillaries and the shank of the electrode was coated with surf wax (Mr Zoggs Sex Wax, CA, USA) to reduce the electrode capacitative transient. For whole-cell recordings the pipette intracellular solution contained (in mM): 131 KCl, 3 MgCl_2_, 1 EGTA-KOH, 5 Na_2_ATP, 5 Hepes-KOH, 10 Na-phosphocreatine (pH was adjusted with 1M KCl to 7.28; osmolality was 294 mmol kg^−1^). In the experiments designed to investigate the effect of extracellular ATP, Na_2_ATP was omitted from the above solution. For cell-attached recordings, the pipette contained (in mM): 140 NaCl, 5.8 KCl, 1.3 CaCl_2_, 0.9 MgCl_2_, 0.7 NaH_2_PO_4_, 5.6 D-glucose, 10 Hepes-NaOH (pH 7.5; 308 mmol kg^−1^). Data acquisition was controlled by pClamp software (RRID:SCR_011323) using Digidata 1320A, 1440A or 1550 boards (Molecular Devices, USA). Recordings were low-pass filtered at 2.5 kHz (8-pole Bessel) and sampled at 5 kHz and stored on computer for off-line analysis (Origin: OriginLab, USA, RRID:SCR_002815). Membrane potentials in whole-cell recordings were corrected for the residual series resistance *R*_s_ after compensation (usually 70-90%) and the liquid junction potential (LJP) of −4 mV measured between electrode and bath solution. The extracellular application of a Ca^2+^-free solution or solutions containing 40 mM KCl, ATP (Tocris Bioscience, UK) or acetylcholine (Sigma-Aldrich, UK) was performed with a multi-barrelled pipette positioned close to the patched cells.

Statistical comparisons of means for electrophysiological experiments and afferent/efferent fibre counting were made by Student’s two-tailed *t* test or for multiple comparisons, analysis of variance (one-way or 2-way ANOVA followed by Bonferroni’s test) were applied. *P*<0.05 was selected as the criterion for statistical significance. Mean values are quoted in text and figures as means ± S.E.M (electrophysiology) and ± S.D. (fibre counting).

### Two-photon confocal Ca^2+^ imaging

For calcium dye loading, acutely dissected preparations were incubated for 40 min at 37°C in DMEM/F12, supplemented with fluo-4 AM (final concentration 10-20 μM; Thermo Fisher Scientific). The incubation medium contained also pluronic F–127 (0.1%, w/v, Sigma Aldrich, UK), and sulfinpyrazone (250 μM) to prevent dye sequestration and secretion. Preparations were then transferred to the microscope stage and perfused with extracellular solution for 20 minutes before imaging to allow for de-esterification.

Ca^2+^ signals were recorded using a two-photon laser-scanning microscope (Bergamo II System B232, Thorlabs Inc., USA) based on a mode-locked laser system operating at 800 nm, 80-MHz pulse repetition rate, <100-fs pulse width (Mai Tai HP DeepSee, Spectra-Physics, USA). Images were formed by a 60x objective, 1.1 NA (LUMFLN60XW, Olympus, Japan) using a GaAsp PMT (Hamamatsu) coupled with a 525/40 bandpass filter (FF02-525/40-25, Semrock). Images were analyzed off-line using custom build software routines written in Python (Python 2.7, Python Software Foundation, RRID:SCR_014795) and ImageJ (NIH) (**Schneider et al., 2012**). Ca^2+^ signals were measured as relative changes of fluorescence emission intensity (ΔF/F_0_). Δ*F* = *F* – *F*_0_, where *F* is fluorescence at time *t* and *F*_0_ is the fluorescence at the onset of the recording

The extracellular application of solutions containing ATP, ryanodine, the P2X antagonist suramin and PPADS (Tocris), the P2Y agonist UTP (Sigma, UK) and the phospholipase C inhibitor U73122 (Tocris Bioscience, UK), was performed using a picospritzer. The pipettes used for local perfusion (diameter 2-4 μm) were pulled from borosilicate glass using a two-step vertical puller (Narishige, Japan). Pressure was kept at a minimum (<3 psi) to avoid triggering mechanically induced calcium signals. Responses in each experimental condition were normalised to control experiments, carried out on the same day under the same imaging and dye-loading conditions.

Each fluorescence recording consisted of 4000 frames taken at 30.3 frames per second from a 125 μm X 125 μm (512 × 512 pixels) region. OHC fluorescence traces were computed as pixel averages from square ROIs (side = 3.7 μm) centred on each OHC. OHCs were classified as either active or inactive using the following algorithm: a) imaging traces were smoothed using a moving average temporal filter of length 3. b) slow Ca^2+^ variations and the exponential decay in fluorescence due to photo-bleaching were removed by subtracting a polynomial fit of order 5 to each trace. De-trended traces were normalized to the maximum value in the recording. c) the noise floor-level was estimated by calculating the power spectral density of the signal using Welch’s method and averaging over the large frequencies (greater than 66% of the Nyquist frequency). d) a spike inference algorithm (*spikes* (**Pnevmatikakis et al., 2016**); module in the *SIMA* python package (**Kaifosh et al., 2014**) was used to estimate the (normalized) spike count *s_i_*. We then calculated the cumulative spike count *S* = Σ*s_i_* for each trace and considered the cell as active (inactive) if *S* was above (below) a predetermined threshold. e) cells that were classified as active (or inactive) and had a maximum signal below (or above) 4 standard deviations were manually sorted. f) The entire dataset was independently reviewed by two experimenters. Cells that had discording classification based on the above criteria (69 out of 2229 at body temperature and 30 out of 5217 at room temperature) were removed from the analysis.

For recording spontaneous activity in the GER, we increased the field of view to a 182 μm X 182 μm region, which was dictated by the ability to detect the full extension of a Ca^2+^ waves in the GER and to maintain a sufficient spatial resolution to resolve the activity of individual OHCs with good signal-to-noise ratio. Under these conditions, the average length of apical coil used for these experiments was 188 ± 4 μm, since some preparations were positioned diagonally in the field of view. Under this recording condition, some large Ca^2+^ waves were underestimated because they were traveling beyond the field of view. Time-series images were corrected for motion using a rigid-body spatial transformation, which does not distort the image (spm12; www.fil.ion.ucl.ac.uk/spm). Recording showing large drifts of the preparation were discarded from the analysis to avoid potential artifacts in the computation of correlation. Calcium waves were manually identified using thresholding and a ROI was drawn around the maximum extension of each multicellular calcium event. Only events that initiated within the field of view of the microscope were considered for this analysis. GER fluorescence traces were computed as ROI pixel averages, and as such they give an indication of the average cytosolic calcium increase in non-sensory cells participating in the propagation of the Ca^2+^ wave. To measure the degree of correlation between OHCs during Ca^2+^ activity in the GER, we computed the average Spearman’s rank correlation coefficient (*r*) between every pair of OHCs in the field of view. We then averaged *r_s_* using Fisher’s z-transformation

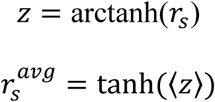

We take 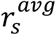 as a measure of the average degree of coordination of the activity of neighbouring OHCs.

To test for the increase in coordinated OHC activity, we used Mann-Whithney U test (one sided) to check whether OHC correlations coefficients during spontaneous Ca^2+^ activity in the GER were significantly (*P*<0.001) greater than those computed during a time window of 400 frames (13.2 seconds) during which no Ca^2+^ waves were observed in the GER.

Photodamage-induced Ca^2+^ waves were triggered by applying high intensity laser pulses using a second mode-locked laser system operating at 716 nm (Mai Tai HP, Spectra-Physics, USA). The laser was merged into the excitation light path using a longpass dichromatic mirror (FF735-Di02, Semrock) and focused on the preparation through the imaging objective (LUMFLN60XW, Olympus, Japan). Two galvanometric mirrors were used to steer the laser beam across the photo-damage area (6.6 × 8.4 um), which typically comprised 1 or 2 non-sensory cells of the GER. The number of repetitions, and thus the total amount of energy delivered, was set to the minimum able to trigger a Ca^2+^ wave (typically 5 repetitions, lasting 165 ms in total).

### Immunofluorescence microscopy

Dissected inner ears from wild-type and Cx30 mice (≥*n* = 4 for each set of experiment) were fixed with 4% paraformaldehyde in phosphate-buffered saline (PBS, pH 7.4) for 5-20 minutes at room temperature. Cochleae were microdissected, rinsed three times for 10 minutes in PBS, and incubated for 1 hour at room temperature in PBS supplemented with 5% normal goat or horse serum and 0.3% Triton X-100. The samples were then incubated overnight at 37°C with the primary antibody in PBS supplemented with 1% of the specific serum. Primary antibodies were: mouse anti-myosin7a (1:1000, DSHB, #138-1), rabbit anti-myosin7a (1:200, Proteus Biosciences, #25-6790), rabbit anti-peripherin (#AB1530, 1:200, Millipore), mouse anti-CtBP2 (1:200, Biosciences, #612044) and rabbit anti-prestin (1:2000, kindly provided by Robert Fettiplace). All primary antibodies were labelled with species appropriate Alexa Fluor secondary antibody for 1 hour at 37°C. Samples were then mounted in Vectashield. The z-stack images were captured with either a LSM 800 with Airyscan (Carl Zeiss) system with GaAsP detectors or with a Nikon A1 confocal microscope. Image stacks were processed with Fiji Image Analysis software.

## Author contribution

F.C., A.H., J-Y.J., S.L.J., D.D.S., W.M. collected and analysed the data. All authors helped with the interpretation of the results and with the writing of the paper. F.C., A.H., M.C.H, C.J.K., D.D.S. and W.M. wrote the paper. WM conceived and coordinated the study.

## Acknowledgements

The authors thank Joerg Striessnig (University of Innsbruck) for providing the *Ca_V_1.3^-/-^* mice; Aubrey Hornak, Andrew Cox and Jemima McCluskey (Baylor University) for their technical assistance with the immunostaining experiments; Michelle Bird (University of Sheffield) for her assistance with the transgenic mouse colonies; Maria Pakendorf (University of Sheffield) for helping with the genotyping. This work was supported by the Wellcome Trust to W.M. (102892), the National Institute on Deafness and Other Communication Disorders to D.D.S. (K18 DC013304) and a 2015–2016 Fulbright Scholar Award to D.D.S. C.J.K. was supported by the MRC (MR/K005561/1).

## References

1. Abe T, Kakehata S, Kitani R, Maruya S, Navaratnam D, Santos-Sacchi J, Shinkawa H (2007) Developmental expression of the outer hair cell motor prestin in the mouse. J Membr Biol 215: 49–56.

2. Ashmore J (2008) Cochlear outer hair cell motility. Physiol Rev 88: 173–210.

3. Beurg M, Safieddine S, Roux I, Bouleau Y, Petit C, Dulon D (2008) Calcium- and otoferlin-dependent exocytosis by immature outer hair cells. J Neurosci 28: 1798–1803.

4. Blankenship AG, Feller MB (2010) Mechanisms underlying spontaneous patterned activity in developing neural circuits. Nat Rev Neurosci 11: 18–29.

5. Bleasdale JE, Fisher SK (1993) Use of U73122 as an inhibitor of phospholipase Ca dependent processes. Neuroprotocols 3: 125–133.

6. Bosher SK, Warren RL (1978) Very low calcium content of cochlear endolymph, an extracellular fluid. Nature 273: 377–378.

7. Boulay AC, del Castillo FJ, Giraudet F, Hamard G, Giaume C, Petit C, Avan P, Cohen-Salmon M (2013) Hearing is normal without connexin30. J Neurosci 33: 430–434.

8. Brown MC (1994) Antidromic responses of single units from the spiral ganglion. J Neurophysiol 71: 1835–1847.

9. Ceriani F, Ciubotaru CD, Bortolozzi M, Mammano F (2016) Design and construction of a cost-effective spinning disk system for live imaging of inner ear tissue. Methods Mol Biol 1427: 223–241.

10. Clause A, Kim G, Sonntag M, Weisz CJ, Vetter DE, Rűbsamen R, Kandler K (2014) The precise temporal pattern of prehearing spontaneous activity is necessary for tonotopic map refinement. Neuron 82: 822–835.

11. Corns LF, Bardhan T, Houston O, Olt J, Holley MC, Masetto S, Johnson SL, Marcotti W (2014) Functional development of hair cells in the mammalian inner ear. Development of Auditory and Vestibular Systems, eds Romand R, Varela-Nieto I (Academic Press), pp. 155–188.

12. Dallos P (1992) The active cochlea. J Neurosci 12: 4575–4585.

13. Delacroix L, Malgrange B (2015) Cochlear afferent innervation development. Hear Res 330: 157–169.

14. Echteler SM (1992) Developmental segregation in the afferent projections to mammalian auditory hair cells. Proc Natl Acad Sci USA 89: 6324–6327.

15. Egan TM, Khakh BS (2004) Contribution of calcium ions to P2X channel responses. J Neurosci 24: 3413–3420.

16. Flores EN, Duggan A, Madathany T, Hogan AK, Márquez FG, Kumar G, Seal RP, Edwards RH, Liberman MC, García-Añoveros J. (2015) A non-canonical pathway from cochlea to brain signals tissue-damaging noise. Curr Biol 25: 606–612.

17. Glowatzki E, Ruppersberg JP, Zenner H-P, Rusch A (1997) Mechanically and ATP-induced currents of mouse outer hair cells are Independent and Differentially Blocked by d-tubocurarine. Neurophamacol 36: 1269–1275.

18. Glowatzki E, Fuchs PA (2000) Cholinergic synaptic inhibition of inner hair cells in the neonatal mammalian cochlea. Science 288: 2366–2368.

19. Guinan JJ Jr. (1996) Physiology of olivocochlear efferents. The Cochlea, eds Dallos P, Popper A, Fay R (Springer, New York), pp 435–502.

20. Hafidi A (1998) Peripherin-like immunoreactivity in type II spiral ganglion cell body and projections. Brain Res 805: 181–190.

21. Housley GD, Marcotti W, Navaratnam D, Yamoah EN (2006) Hair cells--beyond the transducer. J Membr Biol 209: 89–118.

22. Huang LC, Ryan AF, Cockayne DA, Housley GD (2006) Developmentally regulated expression of the P2X3 receptor in the mouse cochlea. Histochem Cell Biol 125: 681–692.

23. Huberman AD, Feller MB, Chapman B (2008) Mechanisms underlying development of visual maps and receptive fields. Annu Rev Neurosci 31: 479–509.

24. Järlebark LE, Housley GD, Thorne PR (2000) Immunohistochemical localization of adenosine 5′-triphosphate-gated ion channel P2X(2) receptor subunits in adult and developing rat cochlea. J Comp Neurol 421: 289–301.

25. Johnson SL, Eckrich T, Kuhn S, Zampini V, Franz C, Ranatunga KM, Roberts TP, Masetto S, Knipper M, Kros CJ, Marcotti W (2011) Position dependent patterning of spontaneous action potentials in immature cochlear inner hair cells. Nat Neurosci 14: 711–717.

26. Johnson SL, Kennedy HJ, Holley MC, Fettiplace R, Marcotti W (2012) The resting transducer current drives spontaneous activity in prehearing mammalian cochlear inner hair cells. J Neurosci 32: 10479–10483.

27. Johnson SL, Ceriani F, Houston O, Polishchuk R, Polishchuk E, Crispino G, Zorzi V, Mammano F, Marcotti W (2017) Connexin-mediated signaling in nonsensory cells is crucial for the development of sensory inner hair cells in the mouse cochlea. J Neurosci 37: 258–268.

28. Jones TA, Leake PA, Snyder RL, Stakhovskaya O, Bonham B (2007) Spontaneous discharge patterns in cochlear spiral ganglion cells before the onset of hearing in cats. J Neurophysiol 98: 1898–1908.

29. Kaifosh P, Zaremba JD, Danielson NB, Losonczy A (2014) SIMA: Python software for analysis of dynamic fluorescence imaging data. Frontiers in Neuroinformatics 8: 77.

30. Katz LC, Shatz CJ (1996) Synaptic activity and the construction of cortical circuits. Science 274: 1133–1138.

31. Katz E, Elgoyhen AB, Gómez-Casati ME, Knipper M, Vetter DE, Fuchs PA, Glowatzki E (2004) Developmental regulation of nicotinic synapses on cochlear inner hair cells. J Neurosci 24: 7814–7820.

32. Kidd EJ, Miller KJ, Sansum AJ, Humphrey PP (1998) Evidence for P2X3 receptors in the developing rat brain. Neuroscience 87: 533–539.

33. King BF, Townsend-Nicholson A (2003) Nucleotide and nucleoside receptors. Tocris Rev 23.

34. Kim G, Kandler K (2003) Elimination and strengthening of glycinergic/GABAergic connections during tonotopic map formation. Nat Neurosci 6: 282–290.

35. Lahne M, Gale JE (2010) Damage-induced cell-cell communication in different cochlear cell types via two distinct ATP-dependent Ca waves. Purinergic Signal 6: 189–200.

36. Lautermann J, ten Cate WJ, Altenhoff P, Grümmer R, Traub O, Frank H, Jahnke K, Winterhager E (1998) Expression of the gap-junction connexins 26 and 30 in the rat cochlea. Cell Tissue Res 294: 415–420.

37. Liberman MC (1980) Efferent synapses in the inner hair cell area of the cat cochlea: an electron microscopic study of serial sections. Hear Res 3: 189–204.

38. Liberman MC, Gao J, He DZ, Wu X, Jia S, Zuo J (2002) Prestin is required for electromotility of the outer hair cell and for the cochlear amplifier. Nature 419: 300–304.

39. Lioudyno M, Hiel H, Kong JH, Katz E, Waldman E, Parameshwaran-Iyer S, Glowatzki E, Fuchs PA (2004) A “synaptoplasmic cistern” mediates rapid inhibition of cochlear hair cells. J Neurosci 24: 11160–11164.

40. Lippe WR (1995) Relationship between frequency of spontaneous bursting and tonotopic position in the developing avian auditory system. Brain Res 703: 205–213.

41. Liu C, Glowatzki E, Fuchs PA (2015) Unmyelinated type II afferent neurons report cochlear damage. Proc Natl Acad Sci USA 112: 14723–14727.

42. Maison SF, Adams JC, Liberman MC (2003) Olivocochlear innervation in the mouse: immunocytochemical maps, crossed versus uncrossed contributions transmitter colocalization. J Comp Neurol 455: 406–416.

43. Marcotti W, Géléoc GS, Lennan GW, Kros CJ (1999) Transient expression of an inwardly rectifying potassium conductance in developing inner and outer hair cells along the mouse cochlea. Pflugers Arch 439: 113–122.

44. Marcotti W, Kros CJ (1999) Developmental expression of the potassium current IK,n contributes to maturation of mouse outer hair cells. J Physiol 520: 653–660.

45. Marcotti W, Johnson SL, Kros CJ (2004) A transiently expressed SK current sustains and modulates action potential activity in immature mouse inner hair cells. J Physiol 560: 691–708.

46. Martinez-Monedero R, Liu C, Weisz C, Vyas P, Fuchs PA, Glowatzki E (2016) GluA2-containing AMPA receptors distinguish ribbon-associated from ribbonless afferent contacts on rat cochlear hair cells. eNeuro 3.

47. Meissner G (1986) Ryanodine activation and inhibition of the Ca2+ release channel of sarcoplasmic reticulum. J Biol Chem 261: 6300–6306.

48. Mellado Lagarde M.M, Cox BC, Fang J, Taylor R, Forge A, Zuo J (2013) Selective ablation of pillar and deiters’ cells severely affects cochlear postnatal development and hearing in mice. J Neurosci 33: 1564–1576.

49. Michna M, Knirsch M, Hoda JC, Muenkner S, Langer P, Platzer J, Striessnig J, Engel J (2003) Cav1.3 (alpha1D) Ca2+ currents in neonatal outer hair cells of mice. J Physiol 553: 747–758.

50. Moody WJ, Bosma MM (2005) Ion channel development, spontaneous activity, and activity-dependent development in nerve and muscle cells. Physiol Rev 85: 883–941.

51. Müller M, von Hünerbein K, Hoidis S, Smolders JW (2005) A physiological place-frequency map of the cochlea in the CBA/J mouse. Hear Res 202: 63–73.

52. Nikolic P, Housley GD, Thorne PR (2003) Expression of the P2X7 receptor subunit of the adenosine 5′-triphosphate-gated ion channel in the developing and adult rat cochlea. Audiol Neurootol 8: 28–37.

53. Oliver D, Plinkert P, Zenner HP, Ruppersberg JP (1997) Sodium current expression during postnatal development of rat outer hair cells. Pflugers Archiv 434: 772–778.

54. Oliver D, Klöcker N, Schuck J, Baukrowitz T, Ruppersberg JP, Fakler B (2000) Gating of Ca2+-activated K+ channels controls fast inhibitory synaptic transmission at auditory outer hair cells. Neuron 26: 595–601.

55. Perkins RE, Morest DK (1975) A study of cochlear innervation patterns in cats and rats with the Golgi method and Nomarkski Optics. J Comp Neurol 163:129–158.

56. Piazza V, Ciubotaru CD, Gale JE, Mammano F (2007) Purinergic signalling and intercellular Ca2+ wave propagation in the organ of Corti. Cell Calcium 41: 77–86.

57. Pnevmatikakis EA, et al. (2016) Simultaneous denoising, deconvolution, and demixing of calcium imaging data. Neuron 89: 285–299.

58. Platzer J, Engel J, Schrott-Fischer A, Stephan K, Bova S, Chen H, Zheng H, Striessnig J (2000) Congenital deafness and sinoatrial node dysfunction in mice lacking class D L-type Ca2+ channels. Cell 102: 89–97.

59. Robertson D (1984) Horseradish peroxidase injection of physiologically characterized afferent and efferent neurones in the guinea pig spiral ganglion. Hear Res 15: 113–121.

60. Rodriguez L, Simeonato E, Scimemi P, Anselmi F, Calì B, Crispino G, Ciubotaru CD, Bortolozzi M, Ramirez FG, Majumder P, Arslan E, De Camilli P, Pozzan T, Mammano F (2012) Reduced phosphatidylinositol 4,5-bisphosphate synthesis impairs inner ear Ca2+ signaling and high-frequency hearing acquisition. Proc Natl Acad Sci USA 109: 14013–14018.

61. Schindelin J, Arganda-Carreras I, Frise E, Kaynig V, Longair M, Pietzsch T, Preibisch S, Rueden C, Saalfeld S, Schmid B, Tinevez JY, White DJ, Hartenstein V, Eliceiri K, Tomancak P, Cardona A (2012) Fiji: an open-source platform for biological-image analysis. Nat Methods 9: 676–682.

62. Shnerson A, Devigne C, Pujol R (1982) Age-related changes in the C57BL/6J mouse cochlea. II. Ultrastructural findings. Dev Brain Res 2: 77–88.

63. Simmons DD (1994) A transient afferent innervation of outer hair cells in the postnatal cochlea. Neuroreport 5: 1309–1312.

64. Simmons DD, Mansdorf NB, Kim JH (1996) Olivocochlear innervation of inner and outer hair cells during postnatal maturation: evidence for a waiting period. J Comp Neurol 370: 551–561.

65. I. Simmons DD, Liberman MC (1998) Afferent innervation of outer hair cells in adult cats: I. Light microscopic analysis of fibers labeled with horseradish peroxidase. J Comp Neurol 270: 132–144.

66. Snyder RL, Leake PA (1997) Topography of spiral ganglion projections to cochlear nucleus during postnatal development in cats. J Comp Neurol 384: 293–311.

67. Spitzer NC (2006) Electrical activity in early neuronal development. Nature 444: 707–712.

68. Teubner B, Michel V, Pesch J, Lautermann J, Cohen-Salmon M, Söhl G, Jahnke K, Winterhager E, Herberhold C, Hardelin JP, Petit C, Willecke K (2003) Connexin30 (Gjb6)-deficiency causes severe hearing impairment and lack of endocochlear potential. Hum Mol Genet 12: 13–21.

69. Tritsch NX, Yi E, Gale JE, Glowatzki E, Bergles DE (2007) The origin of spontaneous activity in the developing auditory system. Nature 450: 50–55.

70. Tritsch NX, Bergles DE (2010) Developmental regulation of spontaneous activity in the Mammalian cochlea. J Neurosci 30: 1539–1550.

71. Wangemann P, Schacht J (1996) Homeostatic mechanisms in the cochlea. The cochlea, eds Dallos P, Popper A, Fay R (Springer, New York), pp 130–185.

72. Weisz CJ, Lehar M, Hiel H, Glowatzki E, Fuchs PA (2012) Synaptic transfer from outer hair cells to type II afferent fibers in the rat cochlea. J Neurosci 32: 9528–9536.

73. Zheng J, Shen W, He DZ, Long KB, Madison LD, Dallos P (2000) Prestin is the motor protein of cochlear outer hair cells. Nature 405: 149–155.

74. Zhao HB, Yu N, Fleming CR (2005) Gap junctional hemichannel-mediated ATP release and hearing controls in the inner ear. Proc Natl Acad Sci USA 102: 18724–18729.

75. Wang HC, et al. (2015) Spontaneous activity of cochlear hair cells triggered by fluid secretion mechanism in adjacent support cells. Cell 163: 1348–1359.

76. Weisz C, Glowatzki E, Fuchs P (2009) The postsynaptic function of type II cochlear afferents. Nature 461: 1126–1129.

77. Zhang-Hooks Y, Agarwal A, Mishina M, Bergles DE (2016) NMDA receptors enhance spontaneous activity and promote neuronal survival in the developing cochlea. Neuron 89: 337–350.

